# Structural basis for allosteric regulation of the proteasome core particle

**DOI:** 10.1101/2024.07.19.604198

**Authors:** Madison Turner, Adwaith B. Uday, Algirdas Velyvis, Enrico Rennella, Natalie Zeytuni, Siavash Vahidi

## Abstract

Intracellular protein degradation is vital across all domains of life^1^. In eukaryotes, the ubiquitin proteasome system performs most non-lysosomal protein degradation and influences numerous cellular processes. Some bacteria, including the human pathogen *Mycobacterium tuberculosis* (*Mtb*), encode a proteasome system that selectively degrades damaged or misfolded proteins crucial for the pathogen’s survival within host macrophages^2–7^. Consequently, the 20S core particle (CP), the central component of the proteasome system, has emerged as a viable target for tuberculosis treatment strategies^2,8–10^. Both eukaryotic and *Mtb* proteasome systems are allosterically regulated^11–13^, yet the specific conformations involved have not been captured in high-resolution structures to date. Here we present the first structure of *Mtb* 20S CP, and indeed any 20S CP, in an inactive state called 20S_OFF_, distinguished from the canonical active state, 20S_ON_, by the conformation of switch helices I and II. The rearrangement of these helices collapses the S1 pocket, effectively inhibiting substrate binding. The switch helices are conserved and regulate the activity of HslV protease, the proteasome’s ancestral enzyme in bacteria, and a diverse family of serine/threonine protein phosphatases. Our results highlight the potential of harnessing allostery to develop therapeutics against the 20S CP in *Mtb* and eukaryotic systems.

## Main

The 20S CP is a barrel-shaped complex composed of four stacked heptameric rings arranged in a conserved α_7_-β_7_-β_7_-α_7_ configuration (Fig. 1a)^4,12,14^. These rings form three interconnected compartments: a pair of antechambers, formed by α_7_-β_7_, that serve as entry points for substrates. Stacked β-rings enclose a degradation chamber where active sites containing the catalytic triad of Thr1, Asp17, and Lys33 proteolyze substrates (Fig. 1b)^6,11,12,14–16^. The activity of the proteasome must be stringently regulated to prevent spurious protein degradation^5^. This is achieved through switching of the conformation of the N-terminal regions of the α_7_ rings. In the absence of stimuli, these “gating” residues extend across the axial pore to cap each end of the 20S CP (Fig. 1c)^15,17^, thereby blocking access to the degradation chamber and mitigating undesired substrate proteolysis. This closed conformation is converted to an open active form through the binding of regulatory particles (RPs)^7,14,17^. The substrate-bound RP docks onto the 20S CP via conserved C-terminal HbYX motifs (Hb: hydrophobic residue, Tyr, X: any residue) that bind between adjacent α-subunits^12,14,19,20^. This triggers conformational changes within the N termini of the α-ring to open the axial gates, thereby allowing for substrate translocation into the degradation chamber by the RP^14^.

**Figure 1.**
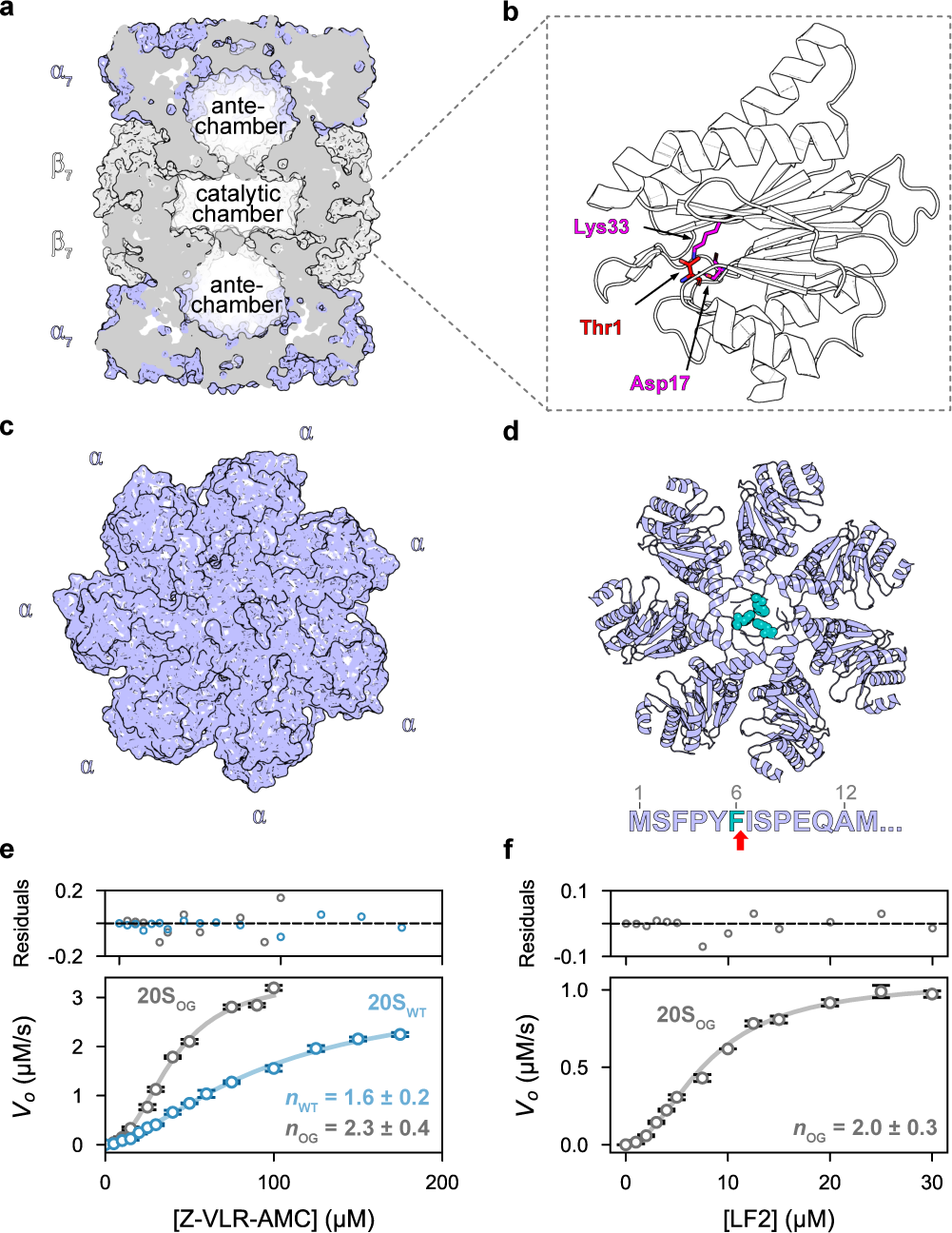
*Mtb* 20S CP displays positive cooperativity upon substrate degradation. (**a**) The 20S CP is a barrel-shaped oligomer with α_7_-β_7_-β_7_-α_7_ architecture surrounding a central catalytic chamber flanked by two antechambers. PDB 3M10; (**b**) Each of the β-subunits is active and contains a catalytic triad consisting of Thr1 (nucleophile, red), Asp17 and Ly33 (magenta); (**c**) Top view of 20S CP_WT_ (left) and 20S CP_OG_ (right) variants. The α-subunits function as gates that cap each end of the proteasome, effectively blocking large substrates from freely diffusing into the catalytic chamber and inhibiting spurious degradation; (**d**) Phe6 (teal) from three of the seven α-subunits block the central pore of the 20S CP when regulatory particles are not bound (PDB 3MKA); (**e**) Functional assays comparing the steady-state kinetics of tripeptide Z-VLR-AMC degradation between the WT and gating-defective (open-gate; OG) 20S CP variants reveal positive cooperativity during substrate hydrolysis. The *n* values from the fits to the Hill model are displayed on the plot; and (**f**) Due to the lack of gating residues, the 20S_OG_ variant was able to degrade the larger, 11-residue peptide LF2, with positive cooperativity. The *n* value from the fit to the Hill model is displayed on the plot.

Insight into the allosteric regulation of *Mtb* proteasome was recently obtained in a study of the interactions between the 20S CP and its RPs, the proteasome accessory factor E (PafE; Rv3780)^11^. Characterization of the binding affinity between a series of *Mtb* 20S CP variants and PafE revealed that the 20S_βT1A_ variant, with a T1A mutation in the β-subunits that impairs catalytic activity (Fig. 1b), and the 20S_OG_ variant, featuring an open gate from the removal of the N-terminal gating residues of the α-subunit (Fig. 1c, d), displayed a higher affinity for PafE than the wildtype (WT) 20S CP^12,13^. This implies that these variants adopt a conformation that is more favourable for RP binding compared to 20S CP_WT_. Yet, these conformations remain elusive in the high-resolution structures obtained to date through X-ray crystallography and electron cryomicroscopy (cryo-EM). Here we reveal near-atomic resolution structures of *Mtb* 20S CP, including a novel inactive state, 20S_OFF_, that interconverts with the canonical active 20S CP state, 20S_ON_. Hydrogen/deuterium exchange mass spectrometry (HDX-MS) measurements highlighted a network of allosterically coupled intra- and inter-ring interactions among α- and β-subunit residues. Mutations targeting these residues allosterically modulate the activity of the 20S CP, illuminating a pathway for the rational design of allosteric inhibitors of this critical enzyme.

### Purification and assembly of *Mtb* 20S CP

We individually expressed and purified α_WT_, α_OG_, β_WT_, and β_T1A_ subunits, and subsequently mixed appropriate pairs of these subunits to assemble three distinct *Mtb* 20S CP variants: 20S_WT_, 20S_βT1A_, and 20S_OG_ (Extended Data Fig. 1). Size exclusion chromatography coupled to multi-angled light scattering (SEC-MALS) showed that all variants had virtually identical elution profiles and estimated molecular weights consistent with the formation of the mature 20S CP (Extended Data Fig. 1e).

### *Mtb* 20S CP displays allosteric enzyme kinetics

As in previous work^6,11,16^, we examined the activity of our 20S CP variants using two substrates. Both of these contain fluorophores whose quantum yield increases upon amide bond cleavage, thereby allowing reaction monitoring via fluorescence. Z-VLR-AMC is a tripeptide that freely diffuses into the degradation chamber of the 20S CP independent of its gating function. Thus, we used this substrate to monitor the intrinsic catalytic activity of the β-subunits. Steady-state kinetic analysis of Z-VLR-AMC hydrolysis by the 20S_WT_ and 20S_OG_ variants produced sigmoidal curves, indicative of positive cooperativity between β-subunits. Fitting of the 20S_WT_ variant profile to the Hill model yielded a Hill coefficient of 1.6 (95% C.I. 1.4 – 1.7), a *K*_0.5_ of 94.5 μM (95% C.I. 80.1 – 111.5), and a *V*_max_ of 3.1 μM s^−1^ (95% C.I. 2.8 – 3.5) (Fig. 1e and Extended Data Fig. 2). Strikingly, the fits for the 20S_OG_ profile yielded an increased Hill coefficient of 2.3 (95% C.I. 1.9 – 2.7), a decreased *K*_0.5_ of 39.1 μM (95% C.I. 35.2 – 43.5), and unchanged *V*_max_ of 3.4 μM s^−1^ (95% C.I. 3.1 – 3.7) as compared to 20S_WT_ (Extended Data Fig. 2). The shift in the Hill coefficient and *K*_0.5_ values between the 20S_WT_ and 20S_OG_ variants points to increased cooperative interactions when the gating residues are removed. Together, these results support the view that the 20S CP is an allosteric system which interconverts between multiple conformations that differ in their activity and affinity for the substrate.

Next, we used a 11-residue substrate (referred to as LF2) that is too large to freely diffuse past the α-gates. Hydrolysis rate of LF2 reports simultaneously on gating and catalytic functions of 20S CP. As anticipated, the 20S_OG_ successfully degraded LF2, notably with a sigmoidal concentration-dependence curve (Fig. 1f). Fitting this activity profile to the Hill model yielded a Hill coefficient of 2.0 (95% C.I. 1.7 – 2.3), a *V*_max_ of 1.1 μM s^−1^ (95% C.I. 1.0 – 1.1) and a *K*_0.5_ of 8.0 μM (95% C.I. 7.1 – 8.9) (Extended Data Fig. 2). By contrast, 20S_WT_ was not active against this larger substrate (data not shown), confirming the expected function of the gates.

Enzyme inhibitors are often employed to stabilize specific protein conformations for detailed structural analysis. Peptidyl boronates, such as ixazomib - a clinically approved human 20S CP inhibitor - represent a class of compounds that competitively inhibit both eukaryotic and prokaryotic 20S proteasomes by mimicking the transition state that the enzyme recognizes during substrate hydrolysis. This renders them ideal for probing the allosteric mechanisms of the *Mtb* 20S CP, as they can stabilize inactive conformations that are otherwise transient and elusive in structural studies^4,21,22^. We found that ixazomib inhibited the peptidase activity of 20S_WT_ against Z-VLR-AMC. Fitting the dose-response curve to a modified Hill model (Eq. 2) yielded an IC_50_ value of 1.1 μM (95% C.I. 0.9 – 1.2) and, notably, a Hill coefficient of 2.1 (95% C.I. 1.6 – 2.6) (Extended Data Fig. 3a, b, c). The potency of this inhibitor enabled its use with our structural tools.

### *Mtb* 20S CP adopts a novel inactive conformation

We used cryo-EM to experimentally determine near atomic-resolution three-dimensional structures of four *Mtb* 20S CP variants: apo 20S_WT_, Ixazomib-bound 20S_WT_, 20S_βT1A_, and 20S_OG_. Subsequent data processing enabled the reconstruction of high-resolution D_7_ symmetry maps refined to global resolutions of 2.7 Å, 2.7 Å, 2.6 Å, and 2.5 Å, respectively (Supplemental Figs. 1-4, Supplemental Table 1). The resolution of all four reconstructed maps was sufficiently high to unambiguously generate molecular models of the 20S CP. The refined models displayed excellent refinement statistics and fit to the corresponding reconstructions (Extended Data Fig. 4, Supplemental Table 1).

All modelled structures adopted the canonical barrel-shaped architecture with the α_7_-β_7_-β_7_-α_7_ arrangement (Supplemental Figs. 1-4). Based on the enforcement of D_7_ symmetry onto the 20S CP maps, we focused our structural analysis on a single representative α-β protomer (Fig. 2a). The density corresponding to the gating residues (αM1-I7) was not resolved in the reconstructed maps of any of the variants, likely due to their conformational flexibility. All four variants have a similar overall structure, as indicated by their low C_α_ RMSD values (0.22 – 0.63 Å) compared to apo 20S_WT_. Notably, 20S_βT1A_ has a slightly higher RMSD value of 0.63 Å compared to the 20S_OG_ variant (RMSD 0.22 Å) and the Ixazomib-bound 20S_WT_ variant (RMSD 0.35 Å), highlighting subtle structural differences between them. A comparison of C_α_ RMSD values calculated separately for the α- and β-subunits showed that conformational changes in the β-subunit primarily account for the observed RMSD deviation in the 20S_βT1A_ variant. The spatial arrangement of the catalytic residues within the 20S CP is essentially unchanged across the four 20S variants (Fig. 2a). Notably, the longitudinal axis of the helix formed by residues βA49-E70, hereafter referred to as switch helix I, shifts by 4 degrees in 20S_βT1A_ compared to all the other 20S variants (Fig. 2a). This results in the displacement of switch helix I by 4.7 Å, which in turn shifts an upstream β-strand and short loop containing residues Ala46, Gly47, Thr48, and Ala49, which form the back of the S1 pocket (Fig. 2b, c). This shift in the loop leads to the S1 pocket closing completely, diminishing its volume from 260 Å^3^ to essentially zero, thereby effectively inhibiting substrate binding. The altered position of switch helix I is stabilized through the formation of a new hydrogen bond between the backbone carbonyl of βG47 and βK33N^ε^. The helix formed by residues βF76-Q96, hereafter referred to as switch helix II, is unchanged in this structure.

**Figure 2.**
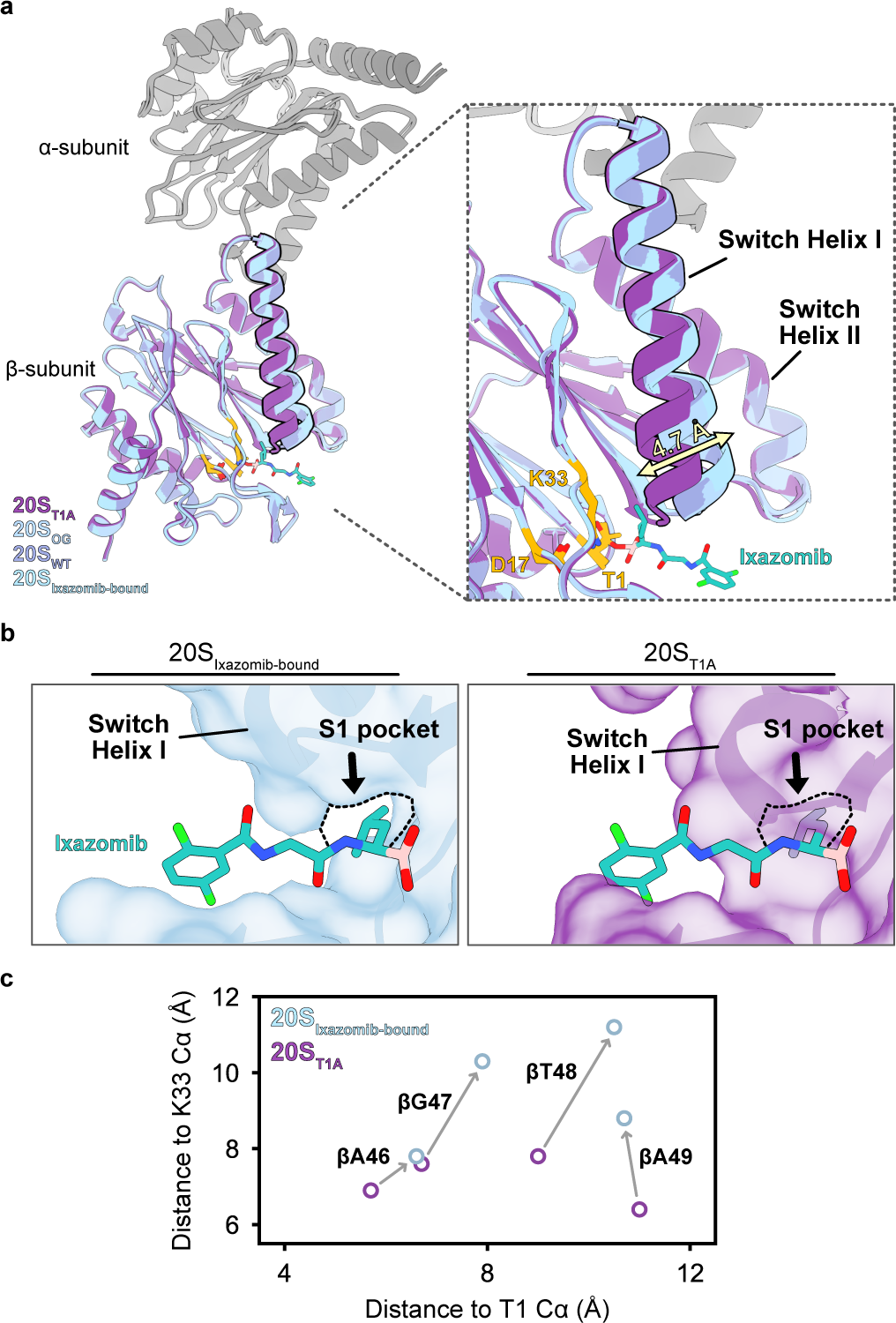
Cryo-EM structural models reveal active and inactive conformations in the proteasome core particle. (**a**) Superimposing one α/β-pair from each cryo-EM structure revealed little difference across the α-subunits from each variant but highlighted a key shift in the β-subunit of 20S_βT1A_ where the Switch helix I shifts by ∼4 degrees and 4.7 Å (measured from C_⍺_ Ala49) compared to the other three variants. (**b**) This movement shifts an upstream loop and β-strand, collapsing the S1 pocket and preventing substrate binding. (**c**) Residues forming the back of the S1 pocket, Ala46, Gly47, Thr48 and Ala49, move up to 3 Å between the 20S_WT_ and 20S_βT1A_ variants when comparing the Cα of the respective residue to the Cα of Thr1 or Lys33 of the catalytic triad.

For the ixazomib-bound 20S_WT_ variant, we observed distinct density features that fit the inhibitor in the active sites (Extended Data Fig. 3d). Consistent with the described inhibition mechanism for this class of proteasome inhibitors^4,21,22^, the boronic acid group of ixazomib is located within 1.5 Å of the catalytic oxygen atom of βThr1 to allow the formation of a covalent bond. In addition, ixazomib binding is mediated by hydrogen bonding with the backbone and surrounding polar side chains (Extended Data Fig. 3d). The leucyl moiety of ixazomib sits in the deep non-polar S1 pocket, where the tert-butyl side chain does not reach. Thus, the binding mode of ixazomib to *Mtb* 20S CP is similar to that observed in its previously determined crystal structure in complex with the human 20S proteasome^3^.

Although the cryo-EM maps of all four variants were refined to similar global resolutions, variations in local resolution are evident when comparing apo 20S_WT_ to the other variants, notably with respect to the densities at the interfaces of the α- and β-rings (Extended Data Fig. 5). Decreased local resolution typically suggests increased conformational variability, indicating localized motion when analyzed across the density map. For example, local resolution in the Ixazomib-bound 20S_WT_ and the 20S_βT1A_ maps is markedly higher and more consistent, especially within the α-rings, compared to resolution in the apo 20S_WT_ and 20S_OG_ maps. This correlates with the loss of structural heterogeneity, particularly in the α-rings, upon ixazomib binding or the introduction of active-site mutation in the β-subunits, again pointing to the presence of an allosteric network spanning across the 20S CP.

### A bidirectional allosteric pathway regulates *Mtb* 20S CP activity

Following structural modeling of the 20S CP variants, we next used hydrogen/deuterium exchange mass spectrometry (HDX-MS) to probe the differences in the conformational dynamics of the same four variants of 20S CP studied using cryo-EM: apo 20S_WT_, Ixazomib-bound 20S_WT_, 20S_βT1A_, and 20S_OG_. HDX-MS is a powerful approach for assessing protein dynamics and structure by measuring the rate at which backbone amide hydrogens exchange with deuterium in D_2_O. Regions of a protein that are less structurally stable exhibit faster exchange rates^23^. HDX-MS is widely used to explore protein-ligand interactions, conformational changes, and other dynamic processes^24^. We performed bottom-up continuous labeling HDX-MS with seven D_2_O exposure times spanning 0-1440 minutes. Peptide mapping yielded a curated list of 78 peptides covering 95% of the α-subunit sequence, and 60 peptides covering 91% of the β-subunit sequence. The average H/D back-exchange was 33% for the α- and β-subunits (Supplemental Fig. 5). We visualized our HDX-MS results as heat maps that display differential deuterium uptake across each 20S CP variant in reference to the apo 20S_WT_ variant. We first focused on the impact of ixazomib binding on the conformational dynamics of 20S_WT_ (Fig. 3a, Extended Data Figs. 6, 7). Twelve β-subunit peptides displayed decreased deuterium uptake. These included the catalytic Asp17 (residues β6-25), the N terminus of switch helix I (residues β49-54), the C terminus of switch helix II (residues βF87-Q98), and the remainder belonged to regions in the immediate vicinity of the catalytic triad and the intra-ring β-β interface. Of note, ixazomib binding also induced changes in the conformational dynamics of the α-subunit. Residues α81-91 at the α-β interface showed a decrease in deuterium uptake at longer D_2_O exposure times. Interestingly, peptides encompassing residues α1-12, which include the gating residues, had elevated deuterium uptake in response to ixazomib binding in the β-subunit. These measurements are supported by multiple partially overlapping peptides.

**Figure 3.**
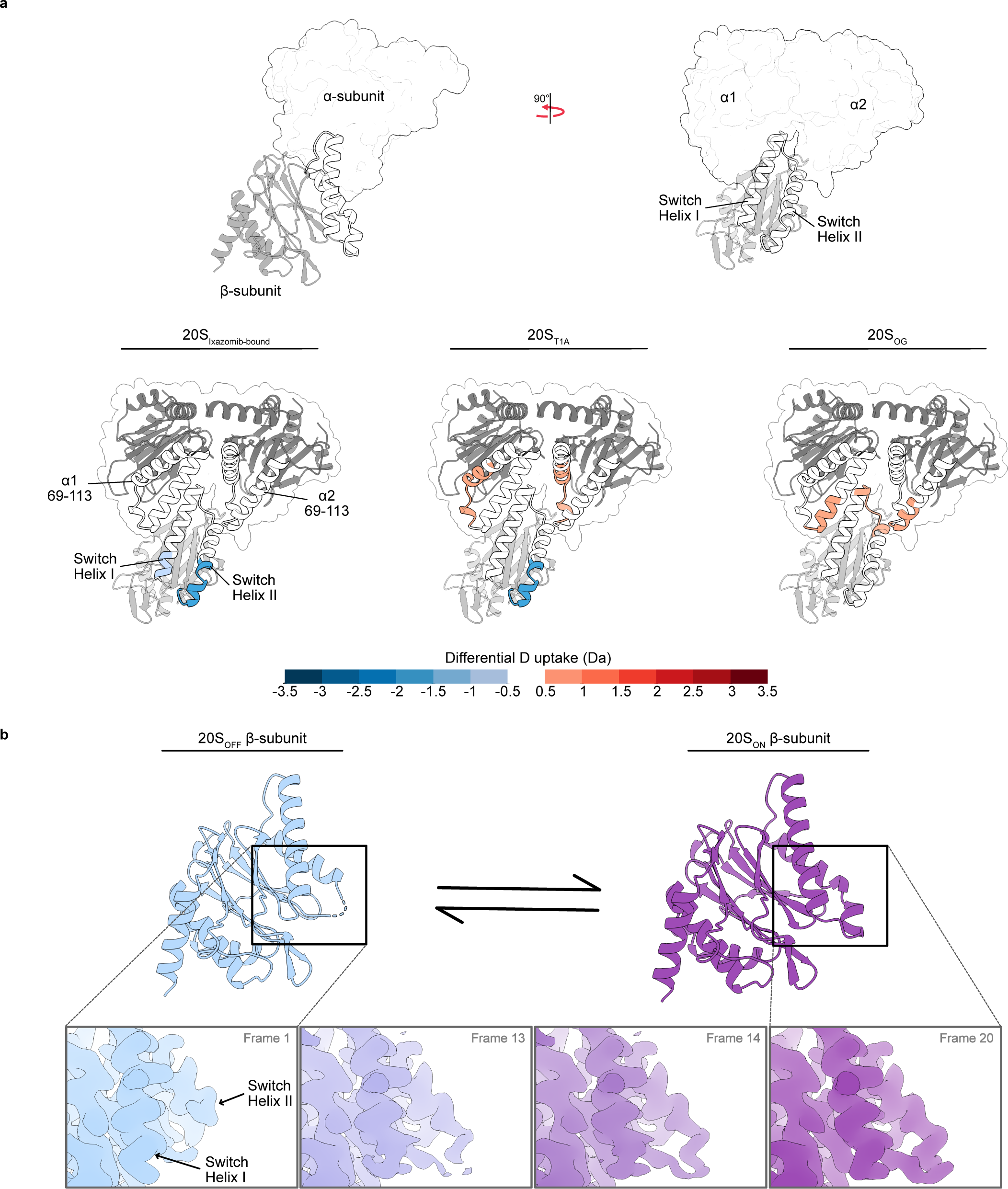
Conformational changes in the switch helices modulate β-subunit activity while also conveying structural information to the α-subunits. (**a**) The switch helices of one β-subunit are in contact with two adjoining α-subunits and represent a likely mechanism for conveying structural information across the α/β-interface. Correspondingly, the two α-subunit helices (residues 69 – 113) proximal to the switch helices show significant differential deuterium uptake compared to the 20S_WT_ variant. Only structural elements of interest are coloured according to the 10-second D_2_O exposure time for each respective state while the remainder of the protein was coloured gray. See Extended Fig. 6, 7 for a complete colour-coded structure. (**b**) Cryo-EM captures the conformational changes in switch helices I and II. Switch helix I changes its position while switch helix II becomes more ordered going from frame 1 to 20. For frames 1 and 20, the refined models are presented above (20S_OFF_ and 20S_ON_, respectively).

We used the 20S_βT1A_ variant to further explore the impact of the active site perturbation on the conformational dynamics of the α-subunit (Fig. 3a, Extended Data Figs. 6, 7). This single-residue substitution in the β-subunit led to differences in deuterium uptake across ten α-peptides and twelve β-peptides. In the β-subunit, peptides containing the catalytic Asp17 and Lys33, and those in contact with the intra-ring β subunits (residues β6-25) showed elevated deuterium uptake relative to apo 20S_WT_. Conversely, the residues in the switch helices I and II (residues β43-50, β49-54, and β87-98) displayed a decrease in deuterium uptake, akin to the pattern observed with ixazomib binding. Likewise, peptides at the α-β interface also demonstrated reduced deuterium uptake, consistent with the changes induced by ixazomib binding. However, the changes span an entire helix in the α-subunit (peptides containing α75-90) rather than just those residues localized to the site closest to the β-ring. Changes in deuterium uptake extended throughout the α-subunit, including the region immediately following the α-gates (residues α19-33) which were less protected compared to apo 20S_WT_ and the structural elements involved in RP binding (Extended Data Fig. 6). Inspection of cryo-EM structures did not show significant differences in the average structure in this region of the 20S CP, highlighting the unique sensitivity of HDX-MS in capturing signatures of conformational flexibility.

Finally, analysis of the differential deuterium uptake of the 20S_OG_ variant relative to 20S_WT_ (Fig. 3a, Extended Data Figs. 6, 7) revealed propagation of conformational changes in the opposite direction. As expected, the structural elements immediately following the site of deletion in the 20S_OG_ variant (residues α13-33) had increased deuterium uptake relative to 20S_WT_. These differences, however, extended throughout the α-subunit and into the β-subunit. In total, twenty-two α-peptides and twenty-five β-peptides had increased deuterium uptake relative to 20S_WT_. We measured elevated deuterium uptake in α-peptides proximal to the α-β interface and in peptides containing βAsp17 and βLys33 of the catalytic triad, and switch helices I and II. Taken together, these HDX-MS results highlight regions that facilitate communication across the α-β interface into the switch helices I and II that in turn modulate the activity of the 20S CP catalytic triad on the β-subunit ∼50 Å away from the gating residues.

### *Mtb* 20S interconverts between active and inactive states on a slow timescale

Representing HDX-MS data in a heatmap format enables global mapping of conformational changes to the structure. We inspected unprocessed HDX-MS data to analyze fine details of deuterium uptake kinetics. Most peptides in the α- and β-subunits have symmetric isotopic distributions across all time points and variants, indicating they adhere to the fast timescale EX2 kinetics. However, several peptides at the α-α, α-β, and β-β interfaces display asymmetric isotopic distributions typically seen in slowly interconverting systems. These include the helix immediately following the gating residues (α13-19), the structural elements that interact with the HbYX motifs at the α-α interface (α41-52, α60-67, α68-74, and α104-116), and other residues at the α-β interface (α75-80, α80-89, and α90-103), including the switch helices I and II (β69-76 and β87-98) (Fig. 3a, Supplemental Fig. 6). For example, the peptide from the C terminus of switch helix II (β87-98) clearly exhibits an asymmetric isotopic distribution in early D_2_O exposure times in the 20S_βT1A_ variant. We globally fit the isotopic distributions of this peptide with the minimum number of Gaussian curves required to achieve robust fits. Each Gaussian represents a distinct conformational state of the 20S. We maintained consistent width and center for each Gaussian component across all variants and time points to minimize the number of fitting parameters, while allowing the amplitudes of the components to vary. This analysis revealed that the populations of the components in the majority of the peptides are anti-correlated while the centre of both distributions increase to higher *m/z* values as a function of D_2_O exposure time, consistent with mixed EX1/EX2 kinetics (Supplemental Fig. 6).

The slowly interconverting conformational states in our HDX-MS data prompted us to perform 3-Dimensional Variability Analysis (3DVA) to uncover both discrete and continuous heterogeneity across our four consensus single-particle cryo-EM maps^25^. 3DVA provides principal components of structural variation. This analysis revealed three distinct motions within our maps. The first motion shows an alternating motion in the α-subunits, as they seemingly compete to occlude the axial pore of the 20S CP (Extended Data Fig. 8 and Supplemental Movie 1). All variants except for 20S_OG_ display this motion, likely due to the absence of the gating residues (α2-6). The second notable component of variability shows contracting and expanding of the β-rings along with rocking motions of the α-rings (Extended Data Fig. 8 and Supplemental Movie 2), a behaviour previously noted in archaeal proteasomes^25^. This motion is present in each 20S variant, except for the ixazomib-bound 20S_WT_ variant, suggesting that inhibitor binding may limit the overall mobility of the 20S CP.

The third component of variability represents the flexible nature of the density corresponding to switch helices I and II and is unique to the 20S_βT1A_ state (Extended Data Fig. 8 and Supplemental Movie 3). Accordingly, we sorted particles from the 20Sβ_T1A_ dataset into 20 intermediate reconstructions along the third principal component from 3DVA and refined them locally to produce maps with resolutions 2.5-2.7 Å (Supplemental Table 2). We subsequently fit our molecular models to the first and last intermediate reconstructed maps (frames 1 and 20, representing the most discrete states in the continuum, respectively) and achieved excellent refinement statistics (Supplemental Table 1). The comparison between the initial and final frames clearly illustrates the transition from the inactive to active conformation of the switch helices. The initial frame closely resembles our modeled inactive consensus structure of 20S_βT1A_ (RMSD 0.44 Å), featuring additional conformational changes in switch helix II (Fig. 3b). The S1 pocket and the conformation of the catalytic residues are unchanged. To our knowledge, this represents the first observation of an inactive conformation of the 20S CP in any species, which we will refer to as 20S_OFF_.

The final frame, designated as 20S_ON_, mirrors the 20S_WT_ variant structure (RMSD 0.49 Å), positioning switch helices I and II in the canonical active conformation captured previously in high-resolution structures (Fig. 3b). Inspection of sequential frames highlights the progressive conformational changes of switch helix I and the decreasing flexibility of switch helix II that accompany the OFF to ON transition. Switch helix I significantly rearranges and moves towards switch helix II. Frames 7 and 8 mark the transition state of the conformational change where both conformations of switch helix I are clearly discernible. Concurrently, residues 92-98 of switch helix II show increasing order, with their density becoming well-defined by frame 20. These findings are further supported by the low local resolution near the tip of switch helix II of the frames noted above and in the consensus map of the 20S_βT1A_ variant compared to the 20S_WT_ variant (Extended Data Fig. 9). Of note, switch helix I shifts by 4.7 Å in the refined atomic model of the 20S_βT1A_ state, while switch helix II undergoes no structural changes in the consensus map as this represents the average state of the particles in the dataset (Fig. 2a).

Overall, our data unveiled 3D structural models of *Mtb* 20S CP under four conditions. We discovered the first conformation of an inactive 20S CP (20S_OFF_), which, compared to the well-known active forms (20S_ON_), exhibits structural modifications in a pair of helices at the α-β interface, termed the switch helices I and II. These helices undergo conformational shifts as the 20S transitions between active and inactive states, culminating in the occlusion of the S1 pocket and effectively modulating substrate accessibility.

### Mutations actuate an allosteric population shift in *Mtb* 20S CP

We used the insights from both cryo-EM and HDX-MS to introduce single-residue mutations in either the α- or β-subunit of the 20S CP to investigate the role of individual residues in the allosteric network (Fig. 4). Although our cryo-EM structures are essentially identical in the α-subunit across all four states, HDX-MS shows allosteric communication in structural elements around the RP-binding site. In the α-subunit, we targeted the RP-binding pocket with two variants, 20S_αS17F_ and 20S_αK52F_, chosen based on a previous study where similar substitutions in the archaeal 20S CP mimicked HbYX binding and induced gate-opening^26^. We hypothesized that these mutations facilitate communication across the α- and β-subunits which is crucial for proteasome activation. For the β-subunit, we targeted Y35 positioned between switch helix I and the active site. In the 20S_βY35F_ variant, the mutation from Tyr to Phe results in the loss of a hydroxyl group essential for H-bonding with the catalytic Lys33 and Val53 on switch helix I. We expected that this substitution would likely weaken the interactions critical for maintaining the active conformation of the switch helices and the S1 pocket of the active site, thereby reducing the activity of the 20S CP. Conversely, in the 20S_βV53Q_ variant would reinstate a β-β intra-ring H-bond found in the archaeal system but absent in *Mtb* 20S CP, reasoning that this would enhance activity by stabilizing the active conformation of the switch helices.

**Figure 4.**
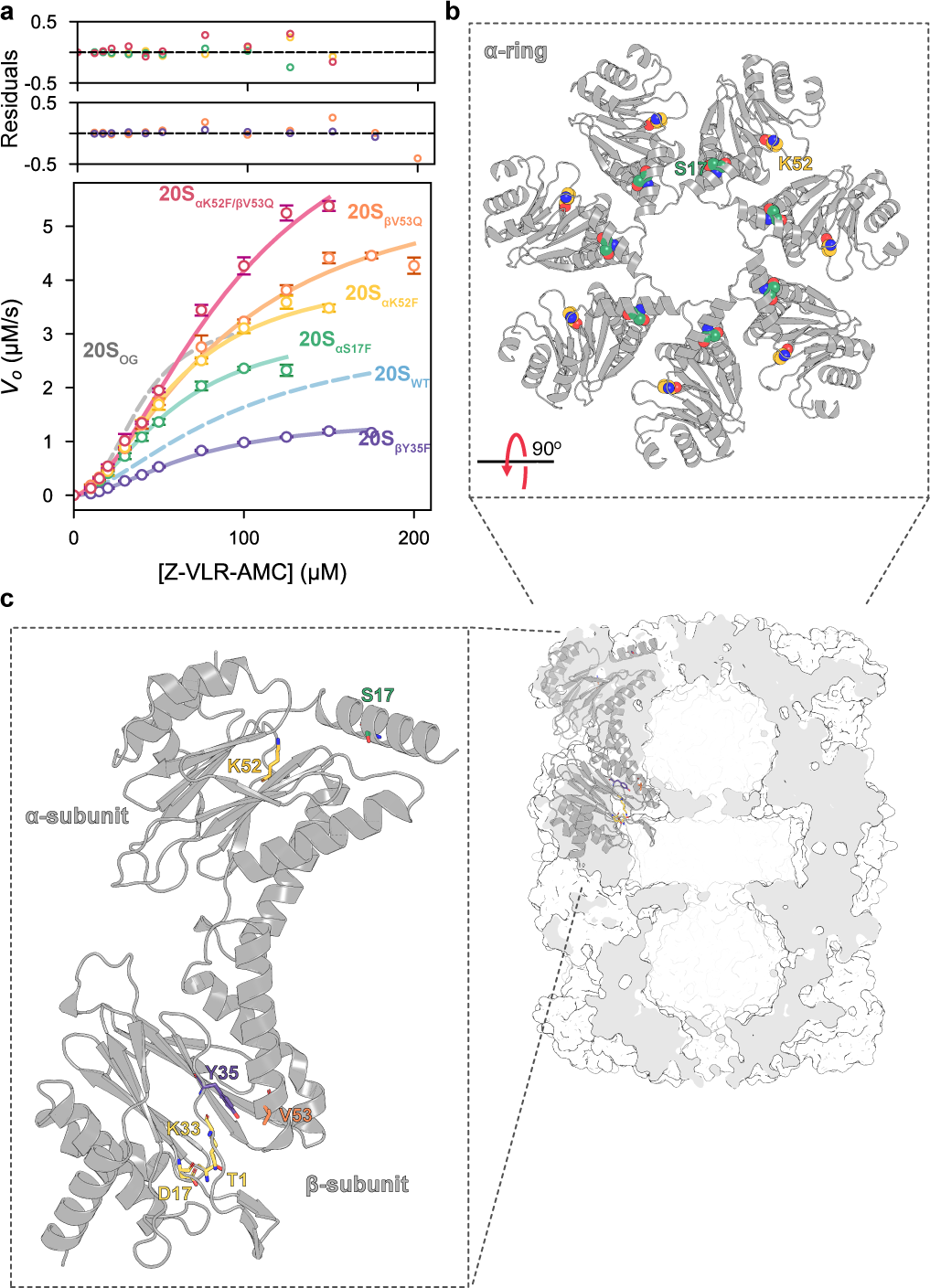
The allosteric equilibrium of *Mtb* 20S CP can be modulated by targeting residues in either the α- or β-subunit. Leveraging insights from both the cryo-EM and HDX-MS studies, a series of point mutations were made to target areas identified as being allosteric hotspots. (**a**) Single residue substitutions to either the α- or β-subunits resulted in modulated catalytic function; (**b**) Residues S17 and K53 of the α-subunits were targeted for mutation due to their role in RP-binding and proteasome activation. In a previous study using the archaeal 20S CP, the substitution of these residues with Phe induced gate-opening; and (**c**) Y35 and V53 of the β-subunits were targeted due to their proximity to the active site and switch helices. The Y35F substitution was made to prevent sidechain-mediated H-bonds that stabilize the active site. V53Q mutation was made to restore a H-bond across intra-ring β-subunits as seen in the archaeal system.

Steady-state kinetic analysis of Z-VLR-AMC hydrolysis by the 20S variants yielded sigmoidal curves, indicative of positive cooperativity. Fitting of the data to the Hill model yielded Hill coefficients ranging from 1.7 to 1.9 (Extended Data Fig. 2). The α-subunit variants, 20S_αS17F_ and 20S_αK52F_, showed *K*_0.5_ values of 57.5 µM (95% C.I. 43.6 – 75.9) and 59.3 µM (95% C.I. 50.5 – 69.8), respectively, while the β-subunit variants, 20S_βY35F_ and 20S_βV53F_, displayed increased *K*_0.5_ values of 68.5 µM (95% C.I. 58.5 – 80.2) and 86.9 µM (95% C.I. 65.8 – 117.0), respectively. Interestingly, while the *V*_max_ of 20S_αS17F_ matched those of the 20S_WT_ and 20S_OG_ variants at 3.2 µM s^−1^ (95% C.I. 26 – 3.9), the 20S_αK52F_ and 20S_βV53Q_ variants exhibited increased *V*_max_ values of 4.2 µM s^−1^ (95% C.I.3.7 – 4.7) and 5.9 µM s^−1^ (95% C.I. 4.9 – 7.2), respectively. In contrast, the 20S_βY35F_ variant had a reduced *V*_max_ of 1.4 µM s^−1^ (95% C.I. 1.3 – 1.6). We combined the most active variants of the α- and β-subunits to determine if the effects of the activating mutations were additive. The 20S_αK52F/βV53Q_ variant maintained unchanged *n* and *K*_0.5_ values compared to the other variants, yet it achieved the highest *V_max_*, reaching 8.0 µM s^−1^ (95% C.I. 6.2 – 10.4). These results underscore the interconnectedness of the α- and β-subunits in 20S CP activity, demonstrating that modifications to either subunit can affect substrate hydrolysis. Notably, none of the variants showed activity against the larger substrate, LF2, which requires gate opening for entry into the 20S chamber, emphasizing the fact that these mutations perturb conformational equilibria within the 20S CP as opposed to the dynamics of the N-terminal gating residues.

### Concluding remarks

Elegant crystallography and cryo-EM studies of *Mtb* 20S CP in isolation^4,15^ and in complex with its RPs^14,20,27^ have revealed many important structural features of this system. Indeed, three of the four states we studied by cryo-EM closely resemble existing crystal structures of the 20S CP. The fourth structure we determined, however, is the first example of *Mtb* 20S CP, and indeed any 20S CP, in an inactive state. This structure showcases a conformational change in a pair of β-subunit helices at the α-β interface, which we have named switch helices I and II, following the convention established by Losick^28^. These switch helices, conserved across all kingdoms of life, regulate the activity of HslV protease^29,30^, the ancestral enzyme of the proteasome ubiquitous in bacteria, and a diverse family of serine/threonine protein phosphatases by orienting the catalytic triad. In the 20S CP, this mechanism does not directly affect the catalytic triad; instead, movement of the switch helices completely obstructs the S1 substrate binding pocket, thereby modulating 20S activity. This regulatory mechanism likely applies to the archaeal^31^ and eukaryotic proteasome, where it is recognized by chaperones that facilitate proteasome maturation^32^. Interestingly, the binding of RPs or mutations at the α-β interface of the archaeal *T. acidophilum* 20S CP modulate substrate specificity, thereby altering the cleavage products^31^. These changes likely result from modifications in the topology of the substrate binding pocket of the 20S CP in that organism. Additionally, comparative analyses of the constitutive proteasome and the immune proteasome suggest that variations in the conformational flexibility of the switch helices underpins the increased activity of the immune proteasome^33,34^. Molecules that selectively bind to the inactive conformation (20S_OFF_) of the proteasome established in this study, rather than the well-known active form (20S_ON_) could be critical to future inhibitor development efforts that focus on allosteric rather than orthosteric sites of the proteasome in bacterial and eukaryotic systems. Our results on *Mtb* proteasome are consistent with those of archaeal and eukaryotic proteasomes that also highlight the importance of dynamics and allosteric control of function^35^. These include allosteric communication between the active sites in the β-rings^36^, between the active sites and the gating residues^26,33,34,37^, between the active sites and the regulatory particles^31,38–42^, and between the α-rings spanning the entire length of the 20S CP^43,44^. Our study extends earlier work on related systems, and brings into focus a view of proteasome regulatory dynamics that is conserved across the three domains of life.

## Acknowledgements

M.T. acknowledges support from a Natural Sciences and Engineering Research Council of Canada Postgraduate Doctoral Scholarship. Financial support was provided by a Canadian Institutes of Health Research Project Grant PJT451412 (S.V.) and a Natural Sciences and Engineering Research Council of Canada Discovery Grant (RGPIN/03031-2022) to N.Z. MS data were recorded at the Mass Spectrometry Facility of the Advanced Analysis Centre, University of Guelph. We thank Dr. Dyanne Brewer (University of Guelph) for assistance with MS measurements. We thank Prof. M. Strauss, K. Basu, and K. Sears at the Facility for Electron Microscopy Research (FEMR) at the McGill University for their assistance with microscope operation, data collection, and computational support. FEMR is supported by the Canadian Foundation for Innovation, Quebec Government, and McGill University. We thank Profs. Lewis Kay (University of Toronto), Matthew Kimber (University of Guelph), and Robert Harkness (University of Guelph) for guidance and helpful discussions.

## Statement of contribution

M.T., E.R., and S.V. initiated the project; M.T., A.B.U., E.R., A.V., N.Z., and S.V. designed research; M.T., A.B.U., E.R., A.V., N.Z., and S.V. performed research; M.T., E.R., A.V., and S.V. contributed new reagents/analytic tools; M.T., A.U.B., N.Z., and S.V. analyzed data; M.T., A.U.B., N.Z., and S.V. wrote the paper; N.Z. supervised the electron cryomicroscopy studies; and S.V. supervised the mass spectrometry and biochemical studies.

## Data availability

Electron microscopy density maps have been deposited in the Electron Microscopy Databank (accession nos. EMD-45494, EMD-45495, EMD-45496, EMD-45498, EMD-45499, EMD-45501, EMD-45532, EMD-45534, EMD-45535, EMD-45537, EMD-45538, EMD-45539, EMD-45540, EMD-45541, EMD-45542, EMD-45547, EMD-45552, EMD-45553, EMD-45556, EMD-45558, EMD-45559, EMD-45560, EMD-45561, EMD-45562). Atomic models have been deposited in the Protein Databank (PDB ID 9CE5, 9CE7, 9CEB, 9CE8, 9CEE, 9CEG). Mass spectrometry data are available from the MassIVE database as entry MSV0000XXXXX.

## Methods

### Plasmids and constructs

Codon-optimized genes encoding the α- (Uniprot: P9WHU1, prcA, Rv2109c) and β-subunit lacking the propeptide (Uniprot: P9WHT9, prcB, Rv2110c) of *Mycobacterium tuberculosis* proteasome core particle (CP) were synthesized and inserted into separate pET24a vectors (Novagen, Madison, WI, USA) at the NcoI and BamHI restriction sites. For the α-subunit constructs, we incorporated a cleavable N-terminal His_6_-SUMO tag. The β-subunit constructs were designed with a C-terminal TEV-SUMO-His_6_ tag to enhance protein solubility during expression and to facilitate the subsequent removal of the tag after purification. Point mutations or deletions were introduced either using Quikchange site-directed mutagenesis or the corresponding genes were synthesized.

### Protein expression, and purification

Three distinct forms of the 20S proteasome CP were expressed, purified, and assembled: 20S_WT_, 20S_αΔ2-6_, and 20S_βT1A_. Unlike previous work that involved the co-expression of the α- and β-subunits^6,15^, we designed a set of constructs along with a purification strategy to express and purify the α- and β-subunits separately. Following purification, these separately-expressed subunits were assembled to form the functional 20S CP. Plasmids were transformed individually into chemically-competent T7 Express *Escherichia coli* BL21(DE3) cells using the heat-shock method and were grown in lysogeny broth (LB) media containing 30 μg/mL kanamycin at 37 °C. When cultures reached an OD_600_ of 0.6, the cells were induced with 1 mM isopropyl β-d-1-thiogalactopyranoside (IPTG) and protein expression was allowed to proceed for 18 hours at 16 °C. Cells were harvested by centrifugation at 4,000 ×*g* for 15 min at 4 °C and resuspended in lysis buffer (300 mM NaCl, 20 mM imidazole and 50 mM Tris-HCl, pH 7.0). The cells were lysed using an Emulsiflex-C3 high-pressure homogenizer (Avestin Inc., Ottawa, Canada) and the resulting homogenate was clarified by centrifugation at 23,700 ×*g* for 50 min at 4 °C. The supernatant was then passed over a Ni^2+^-charged chelating FastFlow™ Sepharose column, which was subsequently washed with 40 mL of lysis buffer, and 30 mL of lysis buffer supplemented with 50 mM imidazole before finally eluting the protein using lysis buffer supplemented with 500 mM imidazole. The α- and β-subunits were subsequently combined and dialyzed against 300 mM NaCl, 1 mM DTT, 50 mM Tris-HCl, pH 7.0 for 18 h at 4 °C in the presence of TEV and Ulp1 proteases to remove the affinity tags and to promote the assembly of the 20S CP complex. The affinity tags and other impurities were then removed by passing the mixture over the IMAC column for a second time. The resulting flow-through was concentrated using a 10 kDa MWCO Amicon Ultra-15 centrifugal filter (Millipore). The fully formed 20S CP was separated from unassembled α- and β-subunits using size-exclusion chromatography (SEC) in 100 mM NaCl and 50 mM NaH_2_PO_4_, pH 7.4 in the running buffer. SDS-PAGE was used at each step to assess protein expression and purity. The final 20S CP concentration was determined spectrophotometrically (guanidinium chloride-denatured protein) using extinction coefficients of 16,390 for the α_WT_-subunit, 14,900 for the α_OG_-subunit, and 20,400 M^−1^cm^−1^ for the β-subunits, obtained from Expasy ProtParam web-based tool (https://web.expasy.org/protparam/).

### SEC-MALS

The oligomeric state of each 20S CP construct was probed using an OMNISEC multi-detector SEC system (Malvern Panalytical, United Kingdom) fitted with an OMNISEC RESOLVE and OMNISEC REVEAL modules. Both the autosampler and column oven were set to 20 °C. Injections of 100 μL with sample concentrations of 2 mg/mL were loaded onto a P3000 Protein SEC column (300×8 mm, Malvern Panalytical) equilibrated in 150 mM NaCl, 50 mM Tris-HCl, pH 7.5 at a flowrate of 1 mL/min. Molecular weight was calculated using light scattering detectors at 90° (right angle light scattering) and 7° (low-angle light scattering). BSA was used as a standard.

### Functional characterization

The peptidase activity of the 20S CP was measured using a pair of substrates: the tripeptide benzyloxycarbonyl-Val-Leu-Arg-7-amino-4-methylcoumarin (Z-VLR-AMC, Genscript) and a 11-residue oligopeptide conjugated to 7-methoxycoumarin-4-acetic acid referred to as LF2^11^ (7-methoxycoumarin4-acetic acid (MCA)-Lys-Lys-Val-Ala-Pro-Tyr-Pro-Met-Glu-(dinitrophenyl)diaminopropionyl-NH2, Genscript). Substrate stocks were prepared in MQ water with 10% DMSO (v/v) at 10-fold the intended final concentration. Each of the 20S proteasome constructs at a concentration of 21.4 nM were incubated with 0 – 175 μM of Z-VLR-AMC and 0 – 30 μM of LF2. At [Z-VLR-AMC] exceeding 175 μM, we observed an inhibition of catalytic activity, which is most likely due to substrate inhibition. Consequently, initial velocities corresponding to [Z-VLR-AMC] concentrations above this threshold were excluded from data fitting. All reactions were performed at 25 °C in 20 mM NaCl, 10 mM NaH_2_PO_4_, pH 7.4 and 1% DMSO (v/v). Reactions were initiated by addition of protein and monitored using a Cary Eclipse fluorometer (Agilent Technologies, Mississauga, Canada). Cleavage of Z-VLR-AMC was monitored using 380 nm and 450 nm excitation and emission wavelengths, respectively, with a bandpass of 5 nm. Degradation of LF2 was monitored at an excitation wavelength of 340 nm and emission wavelength of 405 nm with a bandpass of 5 nm. LF2 was titrated from 0 – 50 μM against 1 μM MCA to correct for the impact of LF2 on MCA fluorescence^45^. The resulting data were fit to a second-order polynomial, and the correction was applied to the measured velocities. Standard curves of AMC or MCA were used to convert relative fluorescence values into the concentration of product formed. Initial velocities were extracted from the first 30 seconds of each reaction using Python scripts written in-house.

Inhibition of the peptidase activity of the 20S WT CP against Z-VLR-AMC was tested using the peptidyl boron-based inhibitor, ixazomib (Selleck Chemicals LLC). Measurements were taken as described above, with a final Z-VLR-AMC concentration of 150 μM and inhibitor concentrations ranging from 0.01 – 10 μM. Both enzyme and substrate stocks were prepared with the appropriate concentration of inhibitor and 1% DMSO (v/v), for a final concentration of 1% DMSO (v/v) in the reaction. Relative activity was then calculated against the uninhibited reaction.

All functional assays were carried out in technical triplicates. Data for both the functional characterization and inhibition studies were fit to the Hill model and visualized using in-house scripts written in Python v3.8.

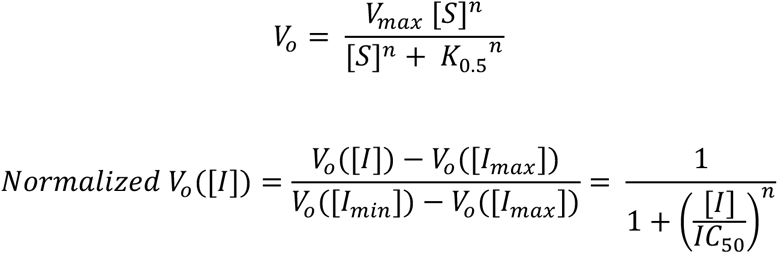

We estimated uncertainties in the fitted *V*_max_, *n*, *K*_0.5_, and apparent IC_50_ parameters using a Monte Carlo approach^46^, wherein we introduced random errors, determined by the root-mean-square deviation between experimental points and those from the optimal-fit model, into the best-fit model to generate 100,000 synthetic datasets. These datasets were then fit as per the experimental data. The values obtained from the Monte Carlo iterations were converted to a histogram, which was subsequently fit to a normal distribution function to yield mean expectation values and standard deviations (*σ*). Log-transformation was used to better estimate the uncertainty of fitted parameters with asymmetric error distributions. Final uncertainties are presented as 2*σ* in the derived values, representing a 95% confidence interval. All relevant scripts are available upon request.

### Cryo-electron microscopy sample preparation and data collection

All 20S CP variants were concentrated to 7 µM, equivalent to 98 µM of each of the α- and β-subunits. For the Ixazomib-bound state, the inhibitor was added to a final concentration of 110 µM in a final DMSO concentration of 0.5% (v/v). Each sample was applied to a holey carbon grid (C-flat CF-2/1-3Cu-T) that had been previously glow-discharged in air at 10 mA for 15 sec. Sample vitrification was performed using a Vitrobot Mark IV (Thermo Fisher Scientific) at 4 °C and 100% humidity. Each sample was incubated on the grid for 3 sec and the grid was blotted for 5 sec with a blot force of 1 before plunging in liquid ethane. Datasets were collected using the SerialEM software^47^ on a Titan Krios microscope (Thermo Fisher Scientific) at the Facility for Electron Microscopy Research at McGill University. Movies were recorded on a Gatan K3 direct electron detector equipped with a Quantum LS imaging filter. The total dose used for each movie was 50 *e^−^*/Å^2^, equally spread over 30 frames. The datasets were collected at a magnification of ×105,000, yielding images with a calibrated pixel size of 0.855 Å. The nominal defocus range used during data collection was between −1.25 µm and −2.75 µm. Data collection parameters for all the datasets are summarized in Supplemental Table 1.

### Cryo-electron microscopy data processing

All cryo-EM data processing was performed using cryoSPARC v4^48^. The movies were corrected for beam-induced motion using patch motion correction, and the CTF parameters were estimated using patch CTF estimation. Micrographs with CTF were fit to a resolution worse than 10 Å and those with unusually high full-frame motion discarded at this stage. The remaining micrographs were used for particle picking using blob picker with a minimum and maximum diameter of 100 Å and 250 Å, respectively. These particles were subjected to two rounds of 2D classification, and the selected classes were used as templates for template-based particle picking. The template-based picked particles were subjected to two rounds of 2D classification for particle curation. The particles from the best classes were then used for *ab-initio* reconstruction and heterogeneous refinement with 5 classes. The classes displaying the 20S CP characteristic structural features were selected and used for high-resolution homogenous refinement. These particles were subjected to reference-based motion correction in cryoSPARC to correct for beam-induced motion at the per-particle level. These motion-corrected particles were subsequently used to obtain the final high-resolution map with D7 symmetry enforced, combined with global and local CTF refinement^49^. Local resolution was estimated for all the maps using cryoSPARC v4. ChimeraX^50^ was used for visualizing the maps and figure preparation. Data processing details for all the datasets are summarized in Supplemental Figs. 1-4.

We used cryoSPARC’s 3D variability analysis (3DVA)^25^ module to probe conformational heterogeneity in our datasets. The particles corresponding to the final refined maps were symmetry expanded to D7 symmetry, and the resulting particles were used as input for the 3DVA with six variability components and a low-pass filter resolution of 3.5 Å. In the case of 20S_βT1A_, the third principal component was subjected to 3DVA intermediate display with a window factor of 0 to obtain 20 non-overlapping classes of particles. This sorted symmetry expanded particles were then used for local refinement to obtain high resolution maps of the intermediate states. The maps corresponding to the two endpoint classes (i.e., frames 1 and 20) were used for atomic modelling.

### Model building and refinement

To build the molecular model for wild-type 20S CP, *Mycobacterium tuberculosis* 20S CP (PDB 8D6V)^27^ was used as an initial template. For all the other datasets, the refined wild-type model was used as the template. The template model was initially docked onto the maps using ChimeraX^50^. For the Ixazomib-bound dataset, the restraints for the threonine hydroxyl-boron bond were generated using JLigand^51^. Model building was performed by an iterative cycle of manual building in Coot^52^ and real space refinement in PHENIX^53^, which significantly improved the model’s geometry and fit in the map. Model validation was also performed on PHENIX^53^, and the model statistics are summarized in Supplemental Table 1.

### HDX-MS

Continuous labeling, bottom-up hydrogen-deuterium exchange mass spectrometry was performed on four states: unbound (apo) 20S_WT_, Ixazomib-bound 20S_WT_, 20S_OG_, and 20S_βT1A_. All solutions used in these experiments contained 100 mM NaCl and 50 mM NaH_2_PO_4_, pH 7.4. D_2_O-based solutions were adjusted to pD 7.4 using the standard electrode correction procedure^54,55^. An initial equilibration step was carried out for each experiment, in which protein stocks were diluted to 2 μM into a H_2_O-based buffer for a minimum of 30 min. 10 μM ixazomib was present in all solutions for the Ixazomib-bound state. HDX was initiated by a ten-fold dilution into D_2_O-based buffer that contained identical additives to that of the equilibrated mixture. HDX was conducted at room temperature with labeling times over 0.167 – 1440 min. Reactions were quenched by acidification to pH_read_ 2.5 by mixing sample aliquots 1:1 (v/v) with 3 M guanidium chloride, 250 mM NaH_2_PO_4_, pH 1.52, and 3 mM n-dodecylphosphocholine^56^ (final concentration of 1.5 M GdnCl, 125 mM NaH_2_PO_4_, and 1.5 mM n-dodecylphosphocholine) and flash frozen in liquid N_2_. Samples were stored at −80 °C prior to analysis.

Liquid handling and reverse-phase separation was performed using a M-Class nanoAcquity UPLC system equipped with HDX technology (Waters, Milford, MA, USA). 5 pmol of sample was digested at 15 °C using a nepenthesin-2 column (Affipro, AP-PC-004, 1 mm × 20 mm, 16.2 µL). The resulting peptides were trapped for 3 min on a BEH C18 (1.7μm, 2.1 mm × 5 mm; Part#: 186003975, Waters) column at a flowrate of 100 μL min^−1^. Peptide separation was achieved on an HSS T3 (1.8 μm, 1.0 × 50 mm; Part#: 186003535, Waters) column using a linear, 8-minute acetonitrile:H_2_O gradient acidified with 0.1% formic acid (acetonitrile ramped from 5 – 35%) at 0 °C and at a flowrate of 100 μL min^−1^. Between each injection, the sample loop was cleaned using 1.5 M guanidine hydrochloride, 4% (v/v) acetonitrile, 0.8% (v/v) formic acid, and 1.5 mM n-dodecylphosphocholine to minimize carry-over.

The ULPC outflow was directed to a quadrupole ion mobility time-of-flight (Q-TOF) Synapt G2-Si mass spectrometer (Waters) fitted with a standard electrospray source operated in positive-ion mode with a capillary voltage of +3 kV. Online calibration of the instrument was achieved by infusing LeuEnk solution (1+, 556.2771 Th) from the LockSpray capillary every 20 seconds at a flow rate of 10 μL min^−1^. Drift time-aligned MS^E^ data-independent acquisition was employed for peptide mapping, as described previously^57^. Data were acquired over the 50 – 2000 *m/z* range with a scan time of 0.4 sec. Fragmentation was induced by linearly increasing the transfer collision energy in alternating scans over 20 – 40 V. Ion mobility separation was controlled manually as described previously^57^. The quadrupole was manually set to dwell at 300 *m/z* to exclude smaller ions. The TOF mass analyzer was operated in resolution mode.

Undeuterated (reference) and deuterated samples were all repeated in technical triplicates. Reference data collected from MS^E^ experiments provided sequence identification using Waters ProteinLynx Global Server (v3.0.3) searched against a database containing the plasmid sequences of both the α- and β-subunits. This database also included variations corresponding to any mutations present in each specific subunit. Peptide filtering parameters were taken from Sørensen et al.^58^. The spectra corresponding to the peptides that satisfied our stringent filtering criteria were manually inspected, and only those with high quality spectra and high signal-to-noise ratio were retained for further analysis. The level of deuterium uptake for any peptide of interest at time *t* is defined as:

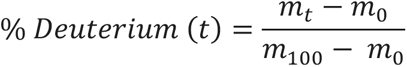

where *m_t_* represents the measured centroid mass of the peptide at time *t*, while *m_0_* and *m_100_* represent the measured centroid masses of the undeuterated and fully deuterated controls, respectively. The undeuterated control underwent identical preparation steps, with the exception that all solution were H_2_O-based. The fully deuterated control was prepared as described by Peterle et al.^59^. Back-exchange was calculated using the maximally deuterated controls.^59^ It should be noted that deletions or mutations were not found to significantly change the predicted *k*_ch_^60^. Data analysis was carried out using DynamX v3.0 (Waters). We used a hybrid significance test model that selected peptides with deuterium uptake differences greater than ± 0.5 Da and that also passed Welch’s *t*-test with α < 0.01, to ascertain significant variations in deuterium uptake^61^.

