## Supplemental Information for "Structural basis for allosteric regulation of the proteasome core particle"

### Supplementary information

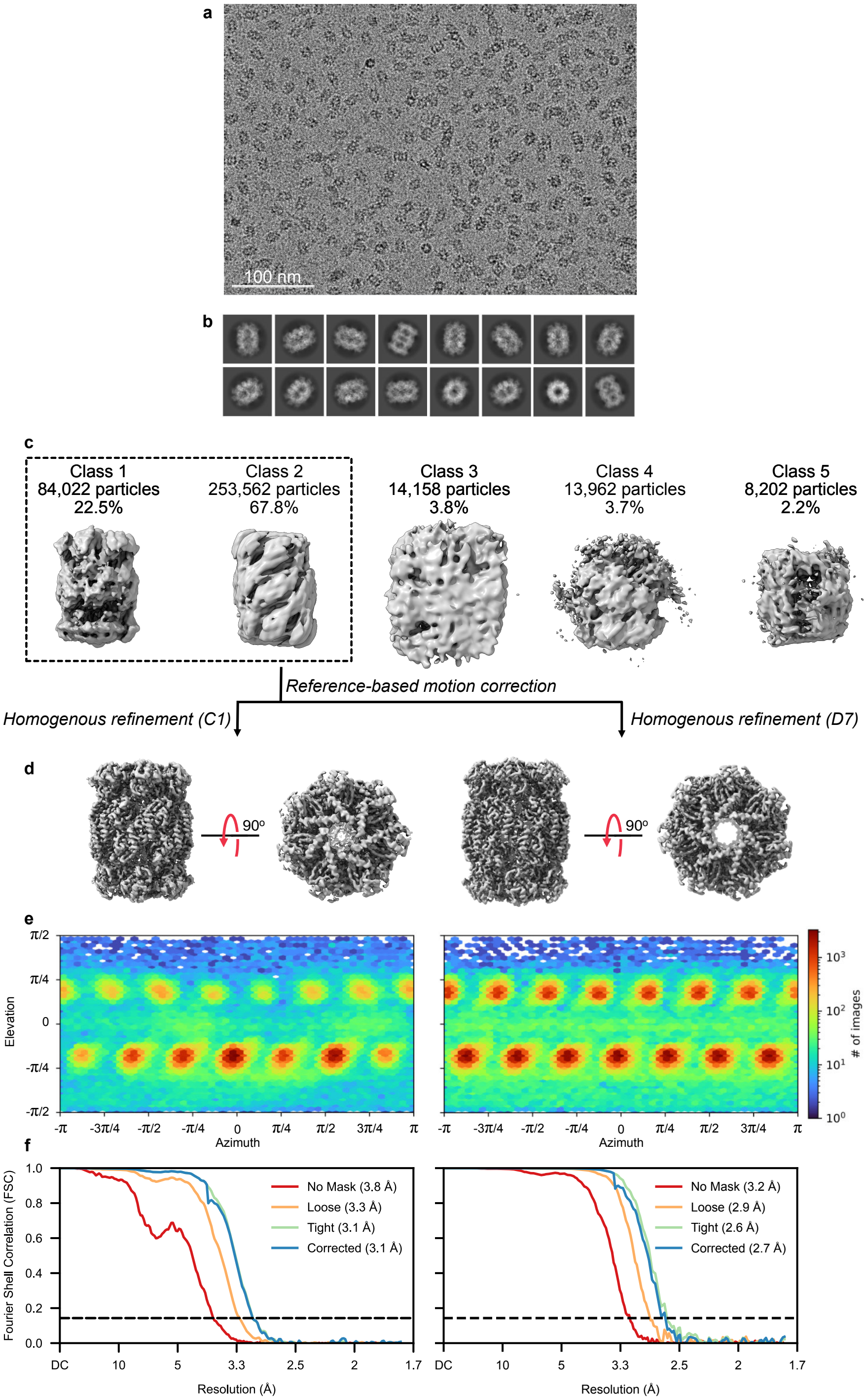


**Supplemental Figure 1. Cryo-EM data processing pipeline for the wild-type 20S core particle.** (a) Representative micrograph (b) Representative 2D classes (c) 3D classes after heterogenous refinement (d) Final consensus maps with C1 (left) and D7 (right) symmetry imposed (e) Viewing direction distribution plots for C1 (left) and D7 (right) refinements (f) Gold-standard Fourier shell correlation plots for C1 (left) and D7 (right) refinements

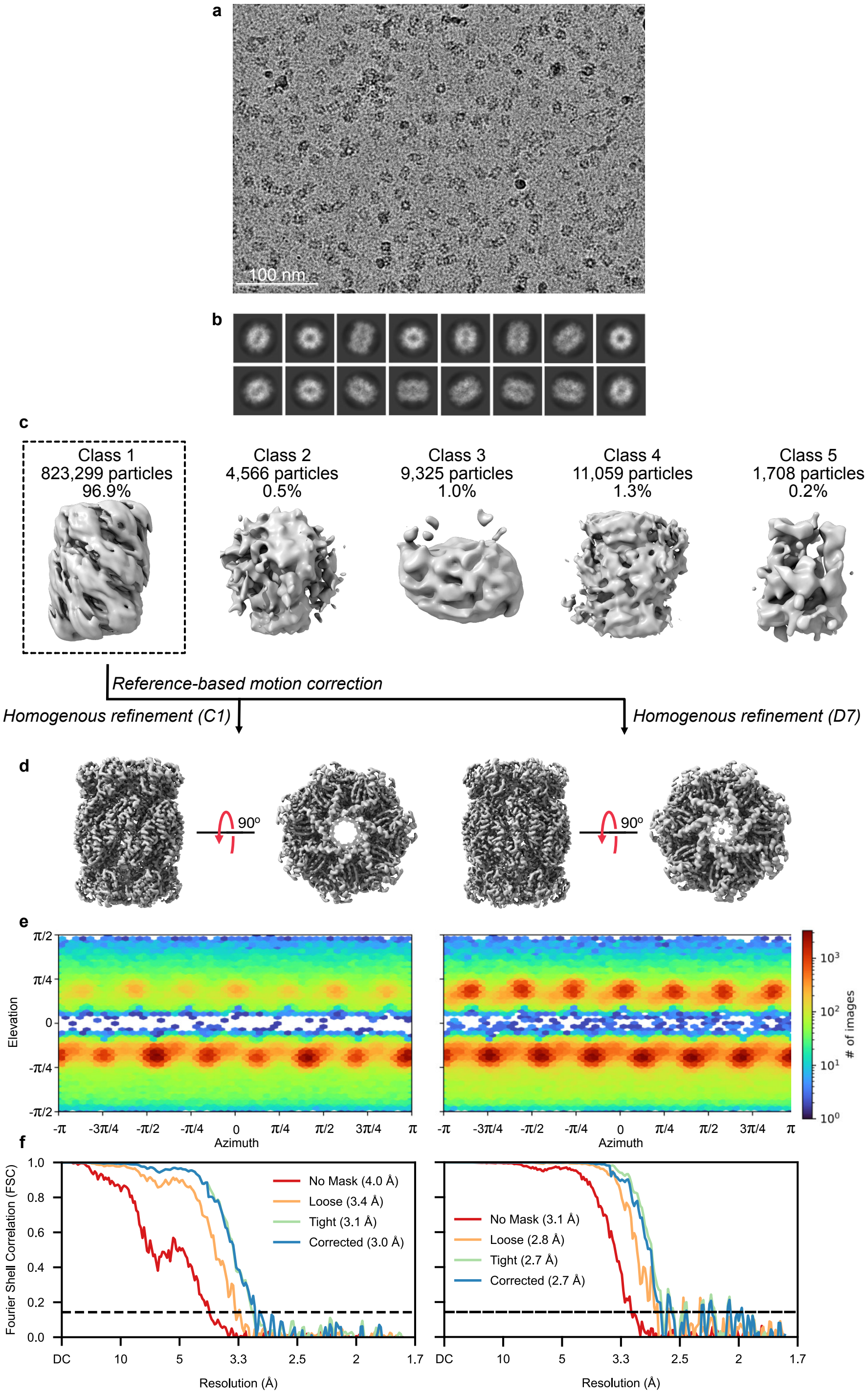


**Supplemental Figure 2. Cryo-EM data processing pipeline for the open-gate 20S core particle.** (a) Representative micrograph (b) Representative 2D classes (c) 3D classes after heterogenous refinement (d) Final consensus maps with C1 (left) and D7 (right) symmetry imposed (e) Viewing direction distribution plots for C1 (left) and D7 (right) refinements (f) Gold-standard Fourier shell correlation plots for C1 (left) and D7 (right) refinements

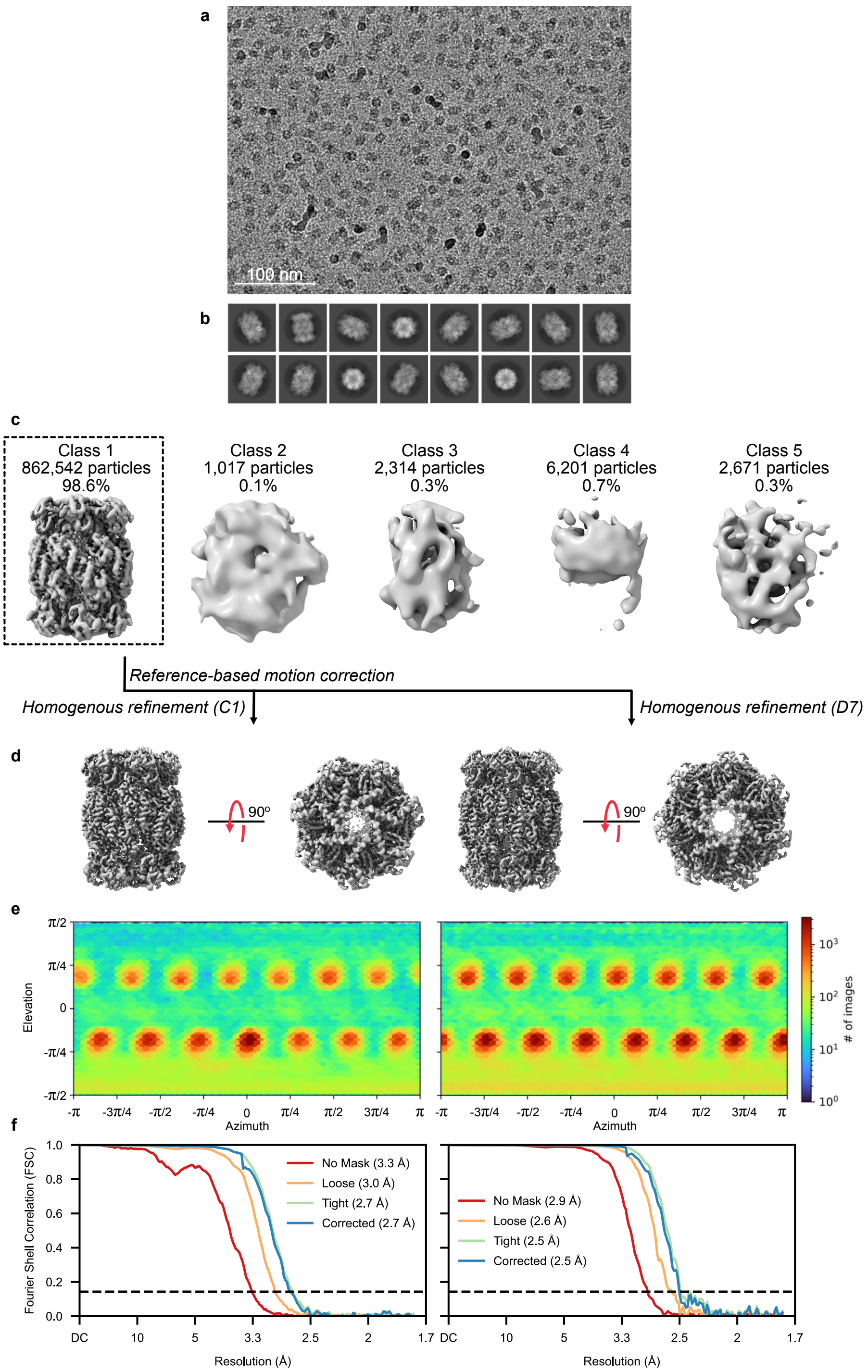


**Supplemental Figure 3. Cryo-EM data processing pipeline for the T1A 20S core particle.** (a) Representative micrograph (b) Representative 2D classes (c) 3D classes after heterogenous refinement (d) Final consensus maps with C1 (left) and D7 (right) symmetry imposed (e) Viewing direction distribution plots for C1 (left) and D7 (right) refinements (f) Gold-standard Fourier shell correlation plots for C1 (left) and D7 (right) refinements

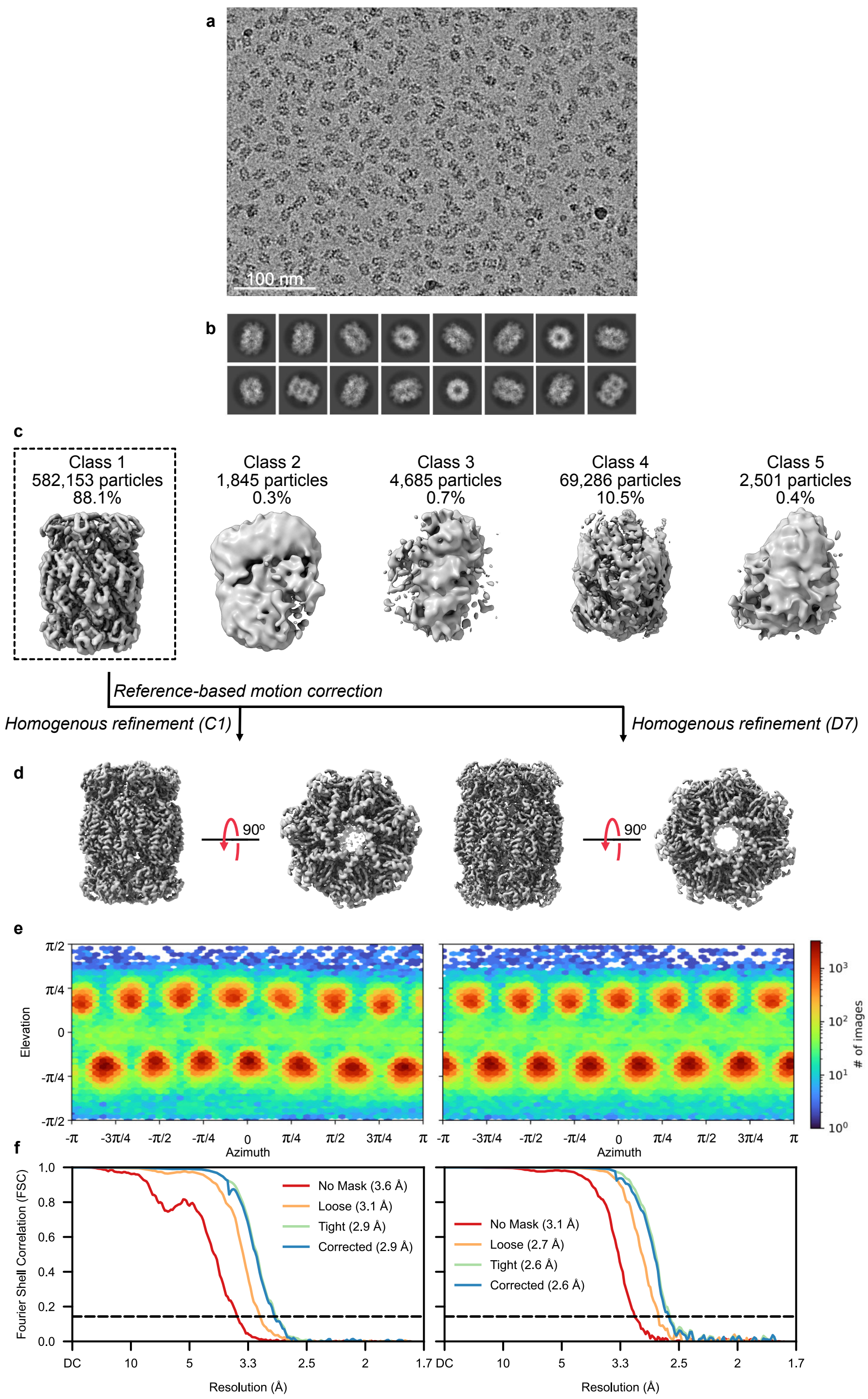


**Supplemental Figure 4. Cryo-EM data processing pipeline for the Ixazomib-bound 20S core particle.** (a) Representative micrograph (b) Representative 2D classes (c) 3D classes after heterogenous refinement (d) Final consensus maps with C1 (left) and D7 (right) symmetry imposed (e) Viewing direction distribution plots for C1 (left) and D7 (right) refinements (f) Gold-standard Fourier shell correlation plots for C1 (left) and D7 (right) refinements

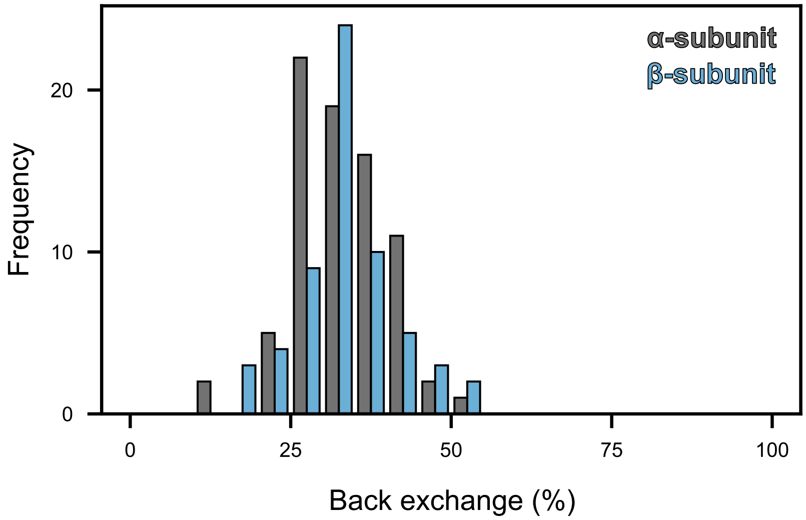


**Supplemental Figure 5. Levels of HDX back-exchange across α- and β-subunit peptides.**

Histogram representing the distribution of α- and β-peptide back-exchange (H to D), expressed as the percentage of deuterium lost from the total number of deuterium incorporated as measured through maximally deuterated samples.

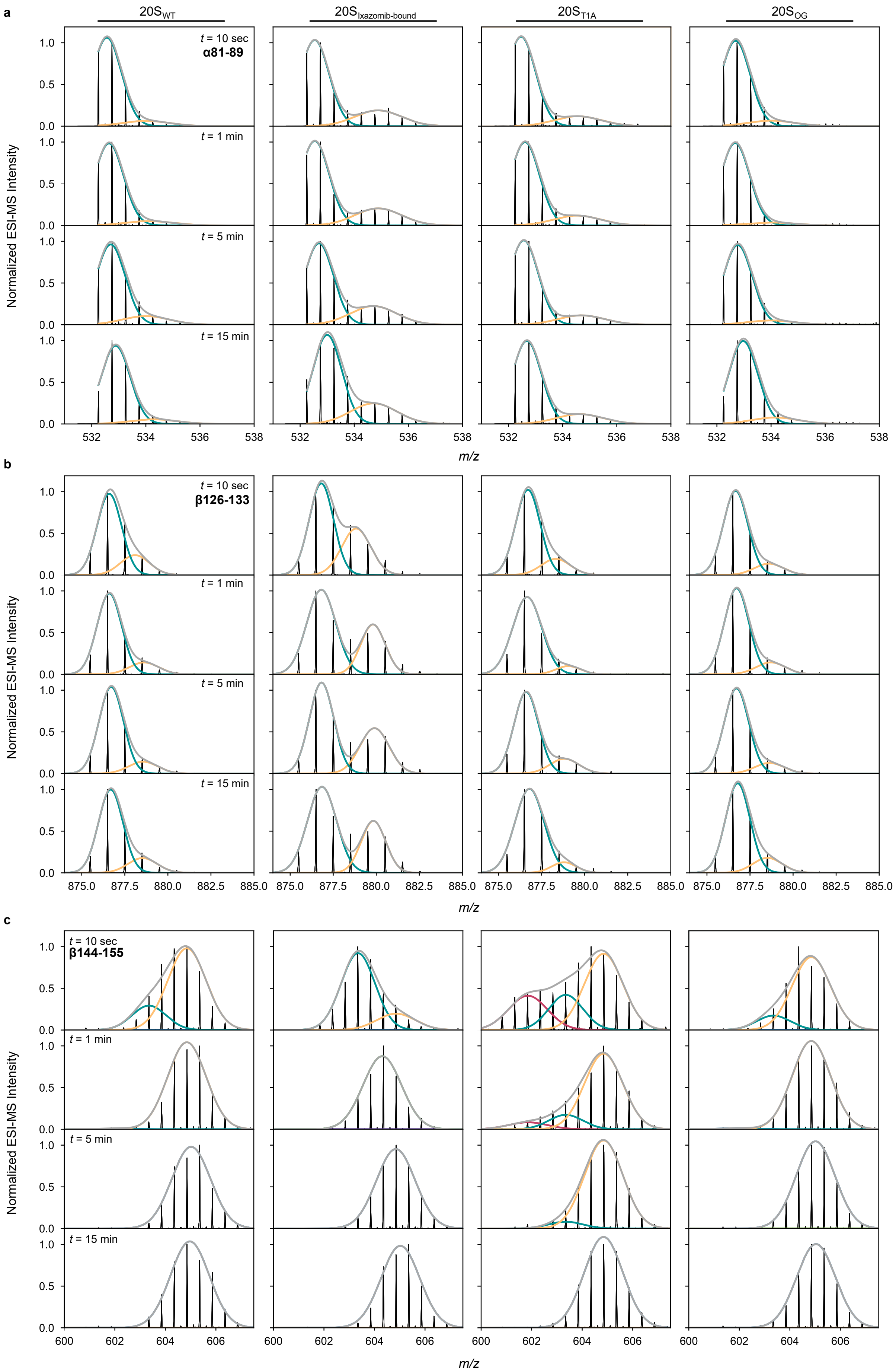
a
b
c
20S_WT
20S_Ixazomib-bound
20S_T1A
20S_OG
t = 10 sec
α81-89
t = 1 min
t = 5 min
t = 15 min
β126-133
β144-155
Normalized ESI-MS Intensity
m/z
0.0
0.5
1.0
532
534
536
538
875.0
877.5
880.0
882.5
885.0
600
602
604
606

**Supplemental Figure 6. HDX-MS isotopic envelopes of 20S$_{βT1A}$ reveal an additional conformation compared to the other variants.**
HDX mass spectra of peptides from the switch helix I in unbound and ixazomib-bound 20S$_{WT}$, 20S$_{βT1A}$, and 20S$_{OG}$ after 10 sec, 1 min, 5 min, and 15 min of $D_2O$ exposure. The grey trace represents the sum of the Gaussian fits.

**Supplemental Movie 1**: 3DVA component 1 for the 20S$_{βT1A}$ variant depicting the alternating motion of the α-subunits.

**Supplemental Movie 2**: 3DVA component 2 for the 20S$_{βT1A}$ variant depicting the overall bending of the 20S CP.

**Supplemental Movie 3**: 3DVA component 3 for the 20S$_{βT1A}$ variant depicting the increased flexibility of the switch helices.

### Supplemental Table 1. Cryo-EM data collection and model building

| | **20SCP WT** | **20SCP OG** | **20SCP T1A (consensus)** | **20SCP-Ixazomib** | **20SCP T1A - OFF state - Frame 1** | **20SCP T1A - ON state - Frame 20** |
|---|---|---|---|---|---|---|
| PDB ID | **9CE5** | **9CE7** | **9CEB** | **9CE8** | **9CEE** | **9CEG** |
| EMDB ID | **EMD-45494** | **EMD-45495** | **EMD-45498** | **EMD-45496** | **EMD-45499** | **EMD-45501** |

**Data collection and processing**

| | | | | | | |
|---|---|---|---|---|---|---|
| Microscope and camera | FEI Titan Krios with Gatan K3 Camera | FEI Titan Krios with Gatan K3 Camera | FEI Titan Krios with Gatan K3 Camera | FEI Titan Krios with Gatan K3 Camera | FEI Titan Krios with Gatan K3 Camera | FEI Titan Krios with Gatan K3 Camera |
| Magnification (x) | 105,000 | 105,000 | 105,000 | 105,000 | 105,000 | 105,000 |
| Voltage (kV) | 300 | 300 | 300 | 300 | 300 | 300 |
| Data acquisition software | Serial EM | Serial EM | Serial EM | Serial EM | Serial EM | Serial EM |
| Electron dose (e-/A2) | 50 | 50 | 50 | 50 | 50 | 50 |
| Defocus range (um) | -1.25 to -2.75 | -1.25 to -2.75 | -1.25 to -2.75 | -1.25 to -2.75 | -1.25 to -2.75 | -1.25 to -2.75 |
| Pixel size (A) | 0.855 | 0.855 | 0.855 | 0.855 | 0.855 | 0.855 |
| Processing software | cryoSPARC v4 | cryoSPARC v4 | cryoSPARC v4 | cryoSPARC v4 | cryoSPARC v4 | cryoSPARC v4 |
| Symmetry imposed | $D_7$ | $D_7$ | $D_7$ | $D_7$ | $C_1$ | $C_1$ |
| Initial particle images (no.) | 437,446 | 1,364,901 | 1,268,434 | 1,238,719 | 11,320,764 | 11,320,764 |
| Final particle images (no.) | 282,632 | 756,031 | 808,626 | 449,574 | 177,137 | 147,239 |
| Map resolution (at FSC=0.143) (A) | 2.66 | 2.72 | 2.5 | 2.61 | 2.89 | 2.86 |

**Refinement**

| | | | | | | |
|---|---|---|---|---|---|---|
| Initial model used | PDB ID: 8D6V | PDB ID: 8D6V | PDB ID: 8D6V | PDB ID: 8D6V | PDB ID: 8D6V | PDB ID: 8D6V |

| | | | | | | |
|---|---|---|---|---|---|---|
| Model resolution - Masked(FSC=0/0.143/0.5) | 2.6/2.6/2.9 | 2.7/2.7/2.7 | 2.4/2.5/2.6 | 2.5/2.6/2.7 | 2.8/2.9/3.1 | 2.7/2.8/3.0 |
| Map sharpening factor (A2) | 130.9 | 134.8 | 130.0 | 131.9 | 93.9 | 89.1 |

**Model Composition**

| | | | | | | |
|---|---|---|---|---|---|---|
| Non-hydrogen atoms | 46144 | 46144 | 46270 | 46466 | 45654 | 46116 |
| Protein residues | 6118 | 6118 | 6118 | 6118 | 6020 | 6118 |
| Ligands | 0 | 0 | 0 | 6V8:14 | 0 | 0 |

**B-factors (A2) (min/max/mean)**

| | | | | | | |
|---|---|---|---|---|---|---|
| Protein | 9.22/121.26/42.42 | 2.11/109.20/32.47 | 0.63/99.24/29.39 | 5.08/110.58/39.59 | 10.01/120.97/50.98 | 19.78/136.20/60.05 |
| Ligand | NA | NA | NA | 32.26/82.69/46.77 | NA | NA |

**r.m.s. deviations**

| | | | | | | |
|---|---|---|---|---|---|---|
| Bond lengths (A) | 0.002 | 0.002 | 0.002 | 0.002 | 0.002 | 0.002 |
| Bond angles (deg) | 0.445 | 0.483 | 0.721 | 0.472 | 0.474 | 0.408 |

**Model Validation**

| | | | | | | |
|---|---|---|---|---|---|---|
| Molprobity score | 1.14 | 1.09 | 1.29 | 1.22 | 1.25 | 1.25 |
| Clash score | 3.48 | 2.8 | 4.49 | 3.61 | 4.85 | 4.59 |
| Poor rotamers (%) | 0 | 0 | 0.3 | 0.3 | 0.61 | 0 |

**Ramachandran plot**

| | | | | | | |
|---|---|---|---|---|---|---|
| Favoured (%) | 98.84 | 97.91 | 97.68 | 97.68 | 98.34 | 97.91 |
| Allowed (%) | 1.16 | 2.09 | 2.32 | 2.32 | 1.66 | 2.09 |
| Disallowed (%) | 0 | 0 | 0 | 0 | 0 | 0 |

**Supplemental Table 2. Cryo-EM data processing parameters for $20S_{\beta T1A}$ intermediate reconstructions.**

Attached as an Excel Spreadsheet.

**Supplemental Table 3. HDX summary**

| | **20S CP WT** | **20S CP OG** | **20S CP T1A** | **20S CP - Ixazomib** |
|---|---|---|---|---|
| HDX reaction details | Final $D_2O$ concentration (v/v) = 90%, $pH_{corr}$ 7.4, RT | Final $D_2O$ concentration (v/v) = 90%, $pH_{corr}$ 7.4, RT | Final $D_2O$ concentration (v/v) = 90%, $pH_{corr}$ 7.4, RT | Final $D_2O$ concentration (v/v) = 90%, $pH_{corr}$ 7.4, RT 10 μM ixazomib ($K_D$ = 1.0 μM); 1% DMSO |
| HDX time course (min) | 0.167, 1, 5, 15, 60, 180, 1440 | | | |
| Undeuterated controls | 3 | | | |
| Back-exchange | α-subunit average back-exchange: 32.8% (range: 13.8 – 52.3 %)<br>β-subunit average back-exchange: 33.3% (range: 16.9 – 53.6 %) | | | |
| Number of peptides | α-subunit: 78 peptides<br>β-subunit: 60 peptides | | | α-subunit: 73 peptides<br>β-subunit: 58 peptides |
| Sequence coverage | α-subunit: 95.2%<br>β-subunit: 91.4% | α-subunit: 97.2%<br>β-subunit: 91.4% | α-subunit: 95.2%<br>β-subunit: 91.4% | α-subunit: 94%<br>β-subunit: 86.7% |
| Average peptide length/redundancy | α-subunit redundancy: 3.55<br>β-subunit redundancy: 3.04 | α-subunit redundancy: 3.55<br>β-subunit redundancy: 3.04 | α-subunit redundancy: 3.55<br>β-subunit redundancy: 3.04 | α-subunit redundancy: 3.23<br>β-subunit redundancy: 3.04 |
| Replicates | 3 technical replicates | | | |
| Repeatability | 0.24 Da | | | |
| Significant differences | 0.5 Da | | | |
