## Extended Data Figures for "Structural basis for allosteric regulation of the proteasome core particle"

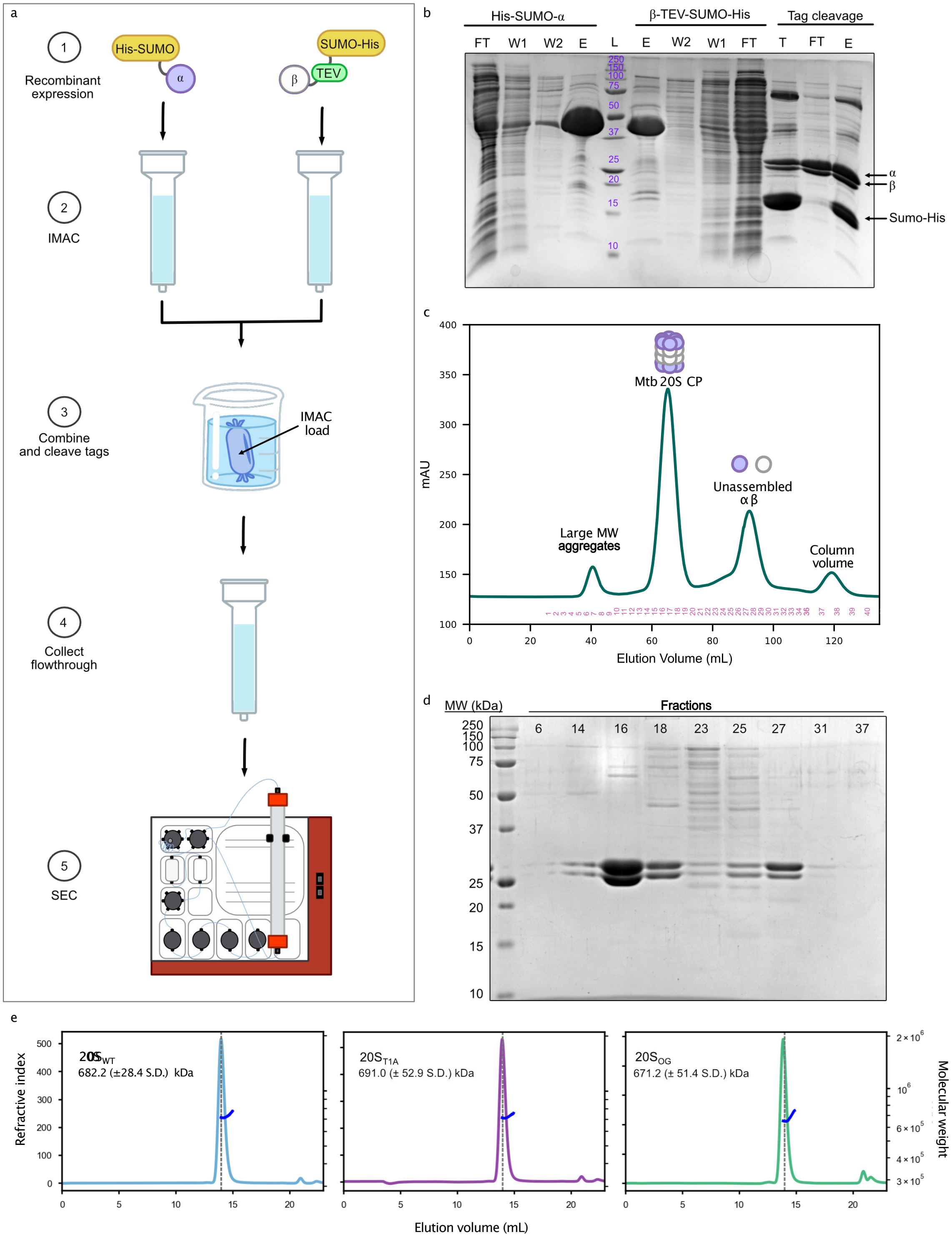


**Extended Data Figure 1. Purification and oligomerization of the mature 20S CP.** (**a**) The purification scheme for 20SP CP starts by isolating individually expressed α- and β-constructs using IMAC (**b**); before combining and purifying assembled core particle using SEC (**c**, **d**); and (**e**) SEC-MALS analysis confirmed the oligomerization state of each variant.

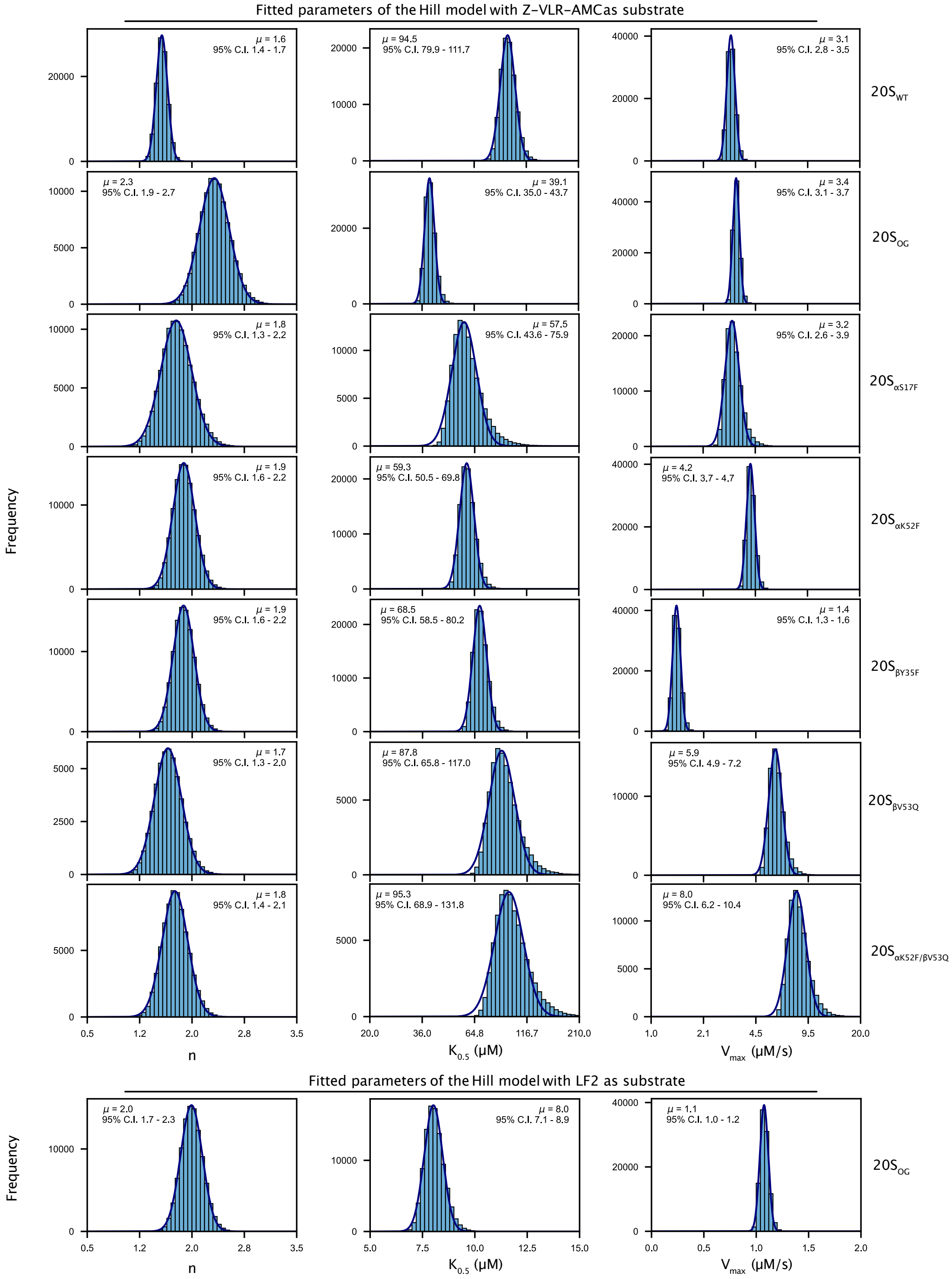


**Extended Data Figure 2. Substrate hydrolysis by the 20S CP CP is well-described by the Hill model**. Histograms representing 100,000 Monte Carlo simulations used to determine the fitted parameters and confidence intervals for *n*, $K_{0.5}$ and $V_{max}$ values of the Hill model fit to the activity data generated using the peptide substrates, Z-VLR-AMC and LF2, for each 20S CP variant.

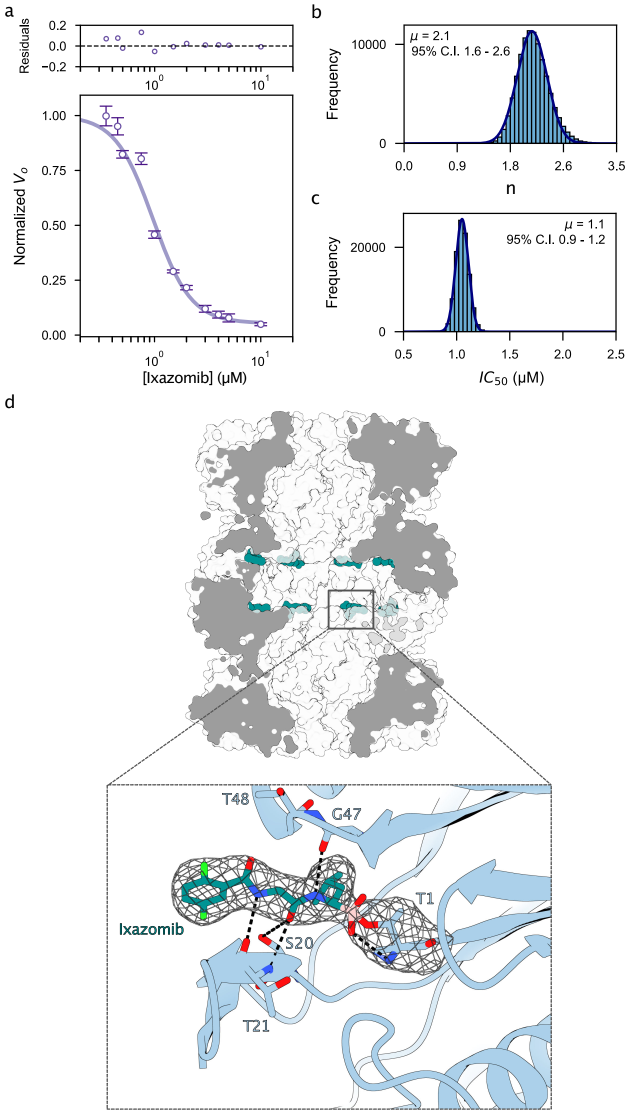


**Extended Data Figure 3. The peptidyl boronate, Ixazomib, functions as a competitive inhibitor against the $20S_{WT}$ CP**. (**a**) Dose-response curve depicting the inhibitory effect of increasing ixazomib concentrations against the hydrolysis of the small tripeptide substrate, Z-VLR-AMC, fitted to the Hill model; (**b**) Histogram representing 100,000 Monte Carlo simulations used to determine the Hill coefficient and respective confidence intervals; (**c**) Histogram representing 100,000 Monte Carlo simulations used to determine the $IC_{50}$ value and respective confidence intervals; and (**d**) Cryo-EM structure of the $20S_{WT}$ CP bound to ixazomib (teal) with density of the small molecule represented as mesh. Density associated with ixazomib was noted in the active site of the CP and displays similar hydrogen bonding patterns to the previously solved structure of the human 20S CP bound to a related peptidyl boronate[3].

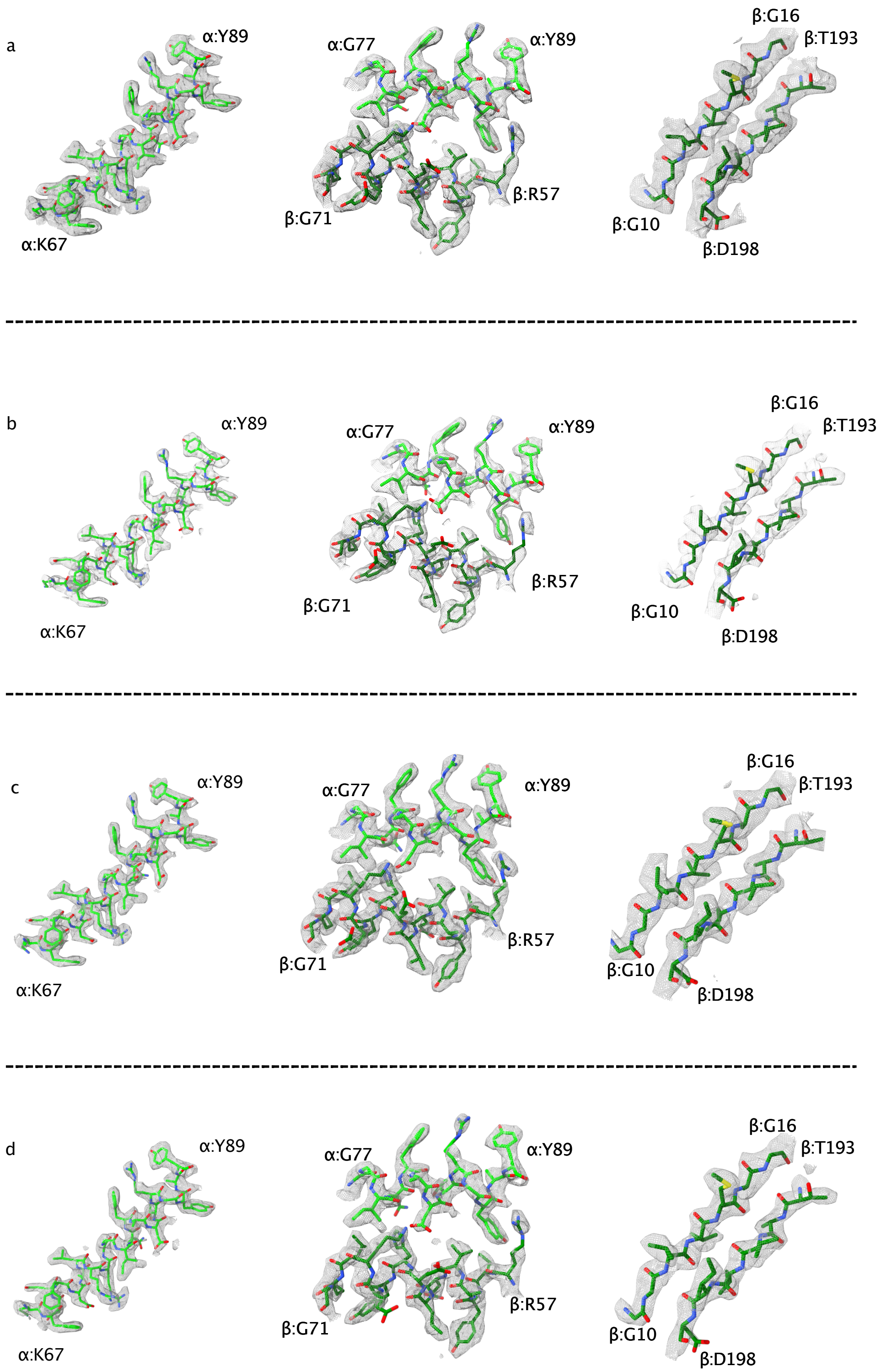


**Extended Data Figure 4. Refined model to consensus map fit for selected regions** (**a**) wild-type 20S core particle (**b**) the open-gate 20S core particle (**c**) T1A 20S core particle (**d**) Ixazomib-bound 20S core particle

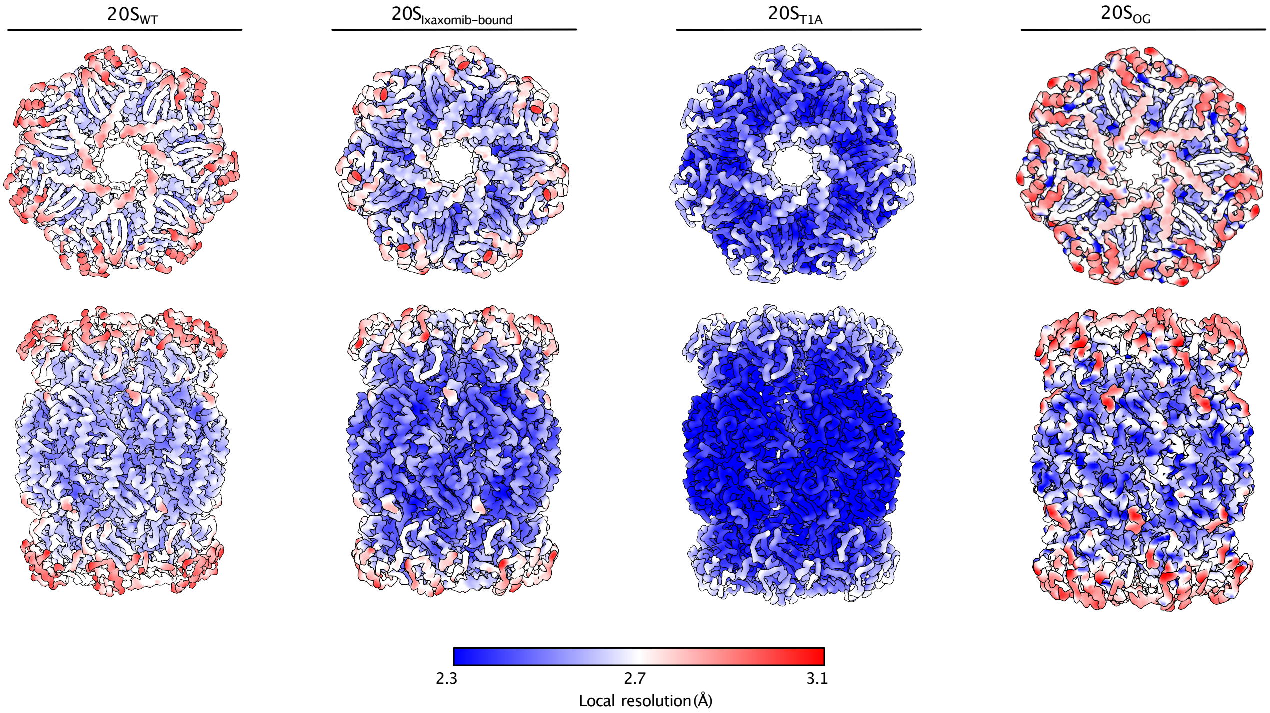


**Extended Data Figure 5. Differences in local resolution across 20S CP variants suggest changes in solution dynamics.** Cryo-EM reconstructed maps of each 20S CP variant coloured according to the local resolution defined by the colour bar. Top (above) and side view (bottom) of each particle map is shown.

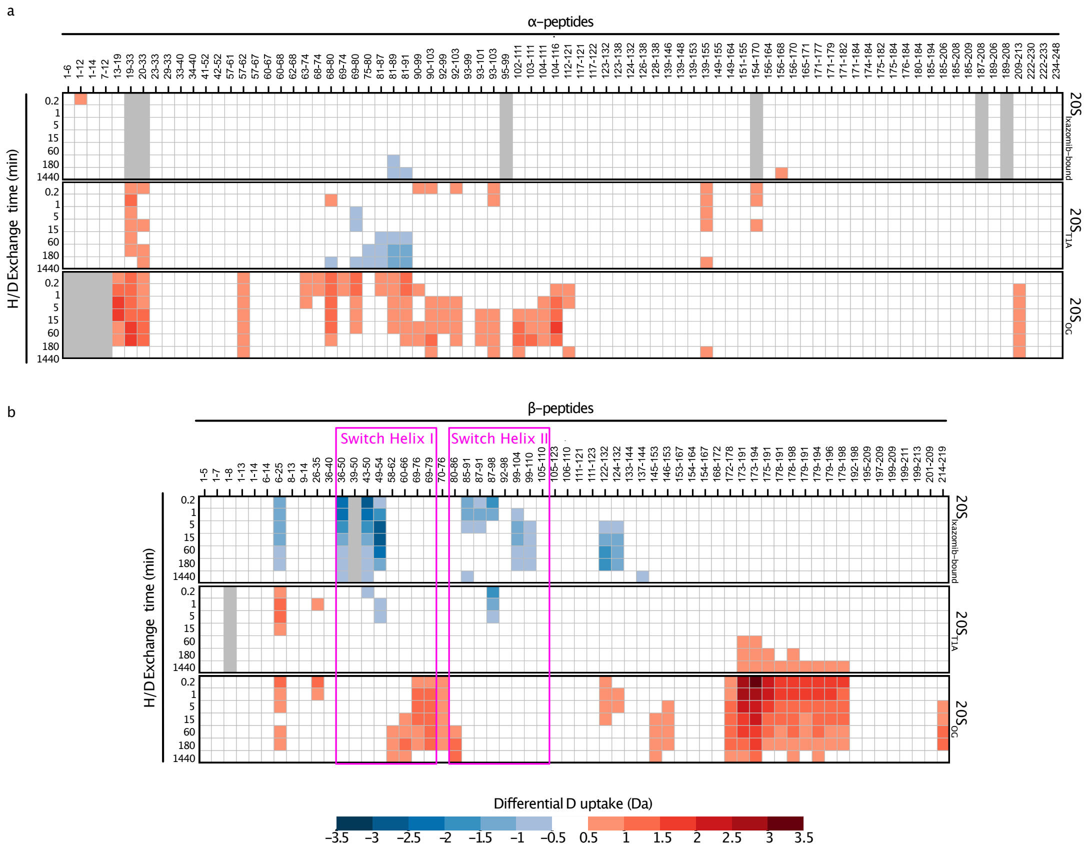


**Extended Data Figure 6. Heat map of relative deuterium uptake highlights the allosteric pathway between the α- or β-subunits of *Mtb* 20S CP.** Changes in deuterium uptake of each state relative to the $20S_{WT}$ variant are depicted for each peptide generated in both the (**a**) α-subunit and the (**b**) β-subunit. Only changes larger than 0.5 Da were considered significant and were coloured according to the colour bar (bottom). The peptides generated for each subunit are listed across the top of the heat map and $D_2O$ exposure times are listed along the side. Gray squares indicate absent data. Peptides associated with the switch helices of the β-subunit are labeled.

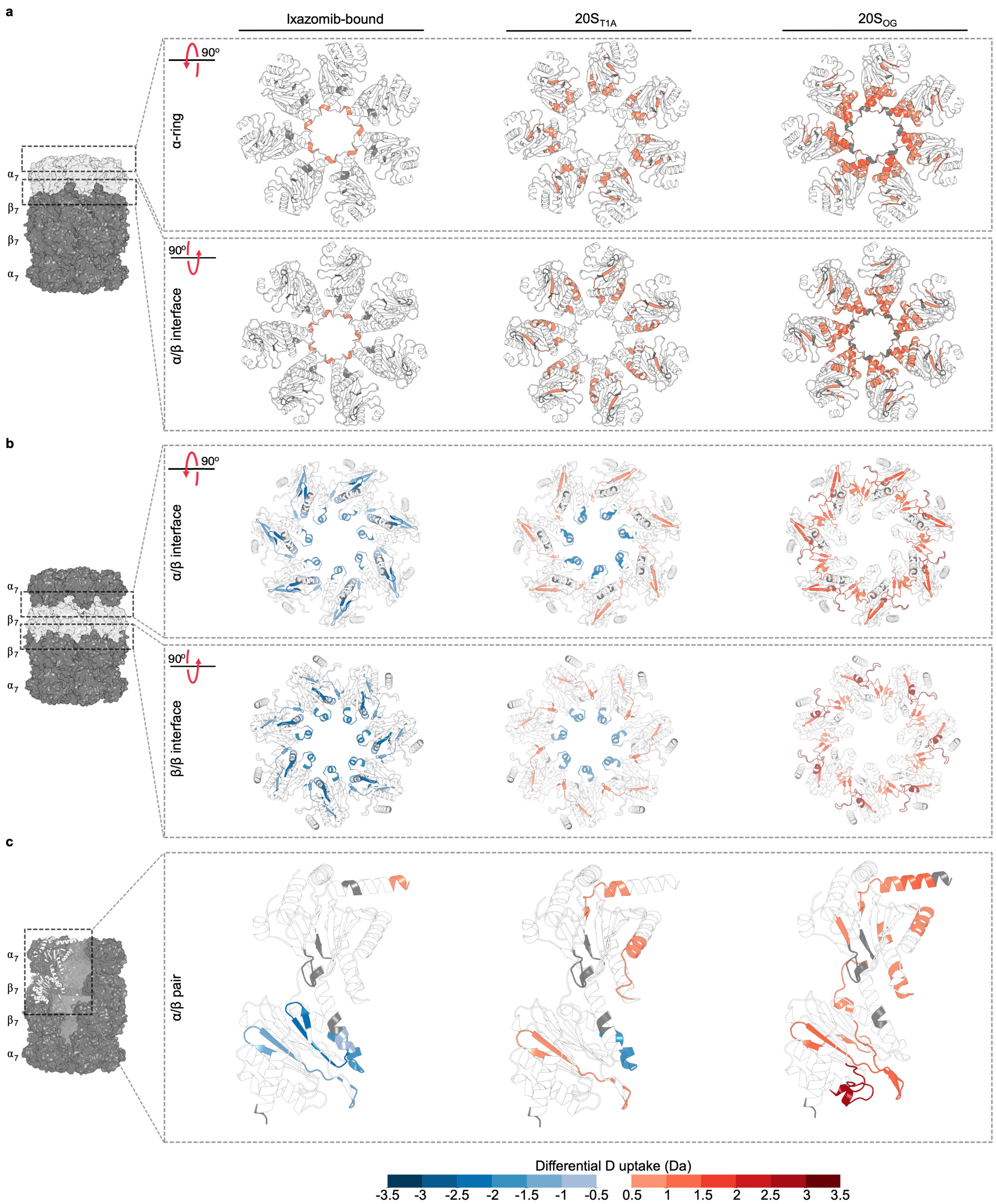


**Extended Data Figure 7. A bidirectional pathway connects the α- and β-subunits of *Mtb* 20S CP.** Relative deuterium uptake after 10 seconds of $D_2O$ exposure colour-coded onto each respective cryo-EM structure highlight regions affected in the (**a**) α-rings, (**b**) β-rings and (**c**) across an α/β pair.

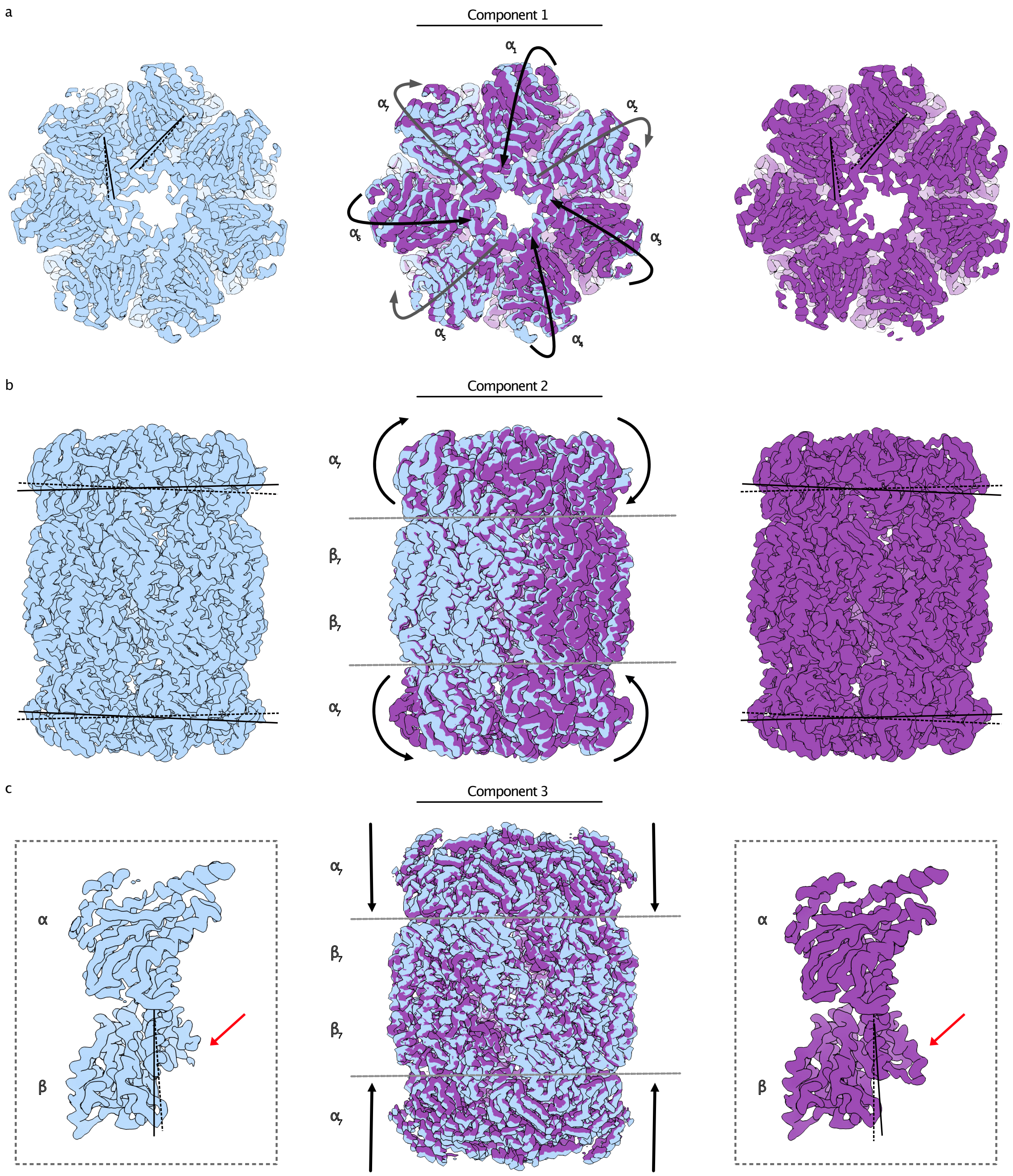


**Extended Data Figure 8. 3D variability analysis of the D7 symmetry expanded particle of the 20S$_{βT1A}$ variant revealed three variability components of the 20S CP.** Maps generated at negative (initial frame, coloured blue) and positive (final frame, coloured purple) latent coordinates along the variability components are overlaid for each component. Solid and dashed lines show the degree of motion between negative and positive frames. Arrows represent the direction of motion. The motions are also shown in Supplemental Movies 1-3 (**a**) The α-subunits compete for occupation of the central pore. (**b**) The α-rings rotate atop the β-rings. (**c**) Compression of the α-rings towards the barrel results in a shift in Switch Helix I and increased density of the Switch Helix II of the β-subunits, highlighted with red arrows, in the 20S$_{βT1A}$ structure.

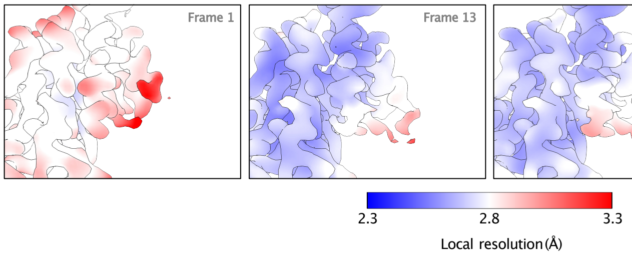


**Extended Data Figure 9. Local resolution of the 3D variability analysis.** Cryo-EM reconstructed maps of frames 1, 13, 14 and 20 of the 3DVA coloured according to the local resolution defined by the colour bar.

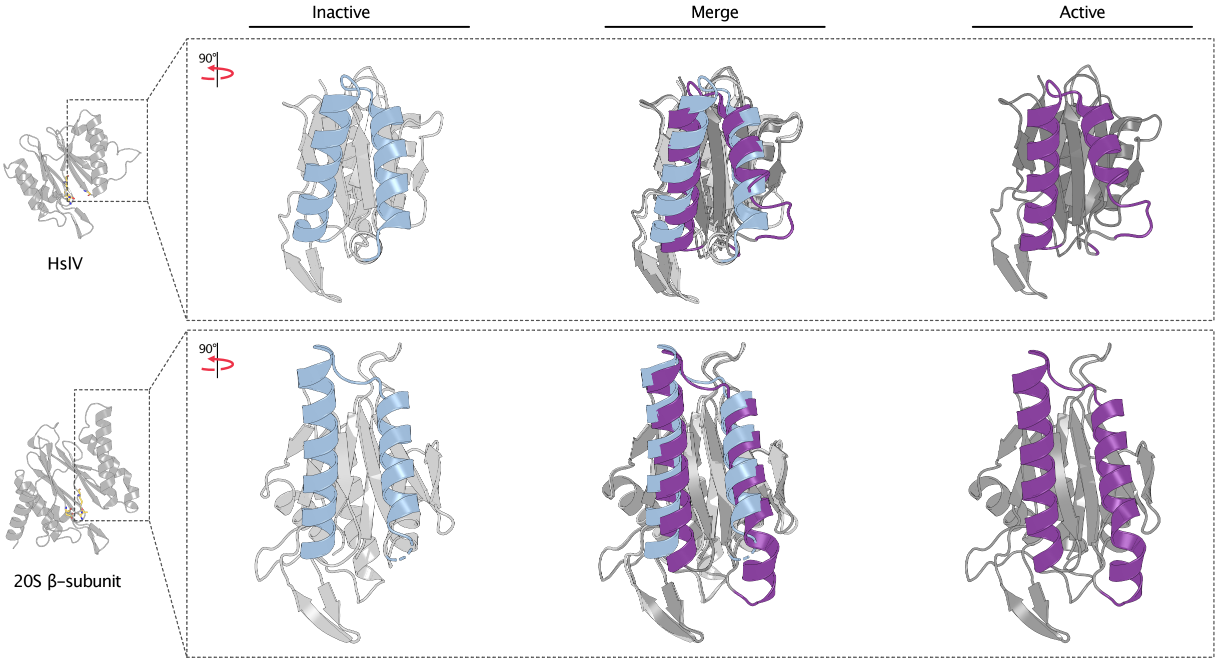


**Extended Data Figure 10. Switch helices are conserved and regulate activity in the primordial enzyme, HslV.** Comparison of the switch helices from the 20S β-subunit to those from HslV, an ATP-dependent protease homologous to the proteasome β-subunit, highlight conserved structural transitions that regulate proteolytic function. The C-terminal region of switch helix II undergoes an order to disordered conformational change between inactive (PDB 1G3K) and active (PDB 1G3I) states of HslV in a manner similar to that seen in the *Mtb* 20S β-subunit.
